# Astrocyte Reactivity by Alcohol Dependence in the Central Amygdala

**DOI:** 10.64898/2026.04.02.716159

**Authors:** Joel G. Hashimoto, Angela E. Gonzalez, Natalie Gorham, Zoe Barbour, Amanda J. Roberts, Le Z. Day, Hermina Nedelescu, Maci Heal, Brett A. Davis, Lucia Carbone, Jon Jacobs, Marisa Roberto, Marina Guizzetti

## Abstract

Astrocytes play essential roles in maintaining brain homeostasis and in contributing to synaptic functions, but, in response to injury, infection, or disease, astrocytes can downregulate their homeostatic and physiological functions while increasing neuroinflammatory responses. The central amygdala (CeA) is important for stress responsivity and the development of alcohol (ethanol) dependence. Using a multi-omics approach in Aldh1l1-EGFP/Rpl10a mice and the chronic intermittent ethanol two-bottle choice (CIE-2BC) model, we have characterized the translational response of CeA astrocytes, as well as the proteomic and phosphoproteomic changes in ethanol dependent, non-dependent, and naïve mice. We identified astrocyte-specific alterations in neuroimmune functions and antioxidant/oxidative stress pathways in ethanol dependent mice as well as cytoskeletal plasticity related pathways in non-dependent mice. Proteomic analysis showed down-regulation of astrocyte physiological functions in dependent animals while phosphoproteomic analysis identified pathways associated with cytoskeleton remodeling in both dependent and non-dependent mice. Reconstructions of astrocyte morphologies demonstrated increased CeA astrocyte complexity in dependent and non-dependent groups compared to naïve mice. The astrocyte-specific activation of neuroimmune and antioxidant pathways, down-regulation of homeostatic functions, alteration in protein phosphorylation-mediated cytoskeleton remodeling, and increased astrocyte morphological complexity demonstrate that ethanol dependence induces astrocyte reactivity in the CeA consistent with both adaptive and maladaptive changes. These findings highlight the role of CeA astrocytes in the progression from alcohol intake to dependence and represent a first step toward identifying astrocyte-specific therapeutic strategies to treat Alcohol Use Disorder (AUD) aimed at potentiating reactive astrocyte adaptive changes and inhibiting maladaptive responses.

## 1. Introduction

Astrocytes play numerous physiological roles in the brain, including maintaining brain homeostasis, modulating synaptic activity, and regulating the functions of the blood brain barrier (MacVicar and Newman, 2015; Mishra, 2017; Farhy-Tselnicker and Allen, 2018; Zhou et al., 2019). Astrocytes also respond to insults to the Central Nervous System (CNS) by becoming reactive, an adaptive response to protect the brain from acute damage (Sofroniew, 2015; Wheeler et al., 2020). Under chronic conditions, however, reactive astrocytes can become maladaptive and induce neuronal damage through the increased production of neuroinflammatory factors and reactive oxygen species (ROS). Maladaptive reactive astrocytes also downregulate astrocyte physiological functions that further contribute to brain dysfunction (Sofroniew, 2020). For these reasons, maladaptive reactive astrocytes are a therapeutic target for several neurodegenerative diseases (Linnerbauer et al., 2020; Lee et al., 2022). Reactive astrocytes exhibit stimulus-, disease-, and time-dependent molecular, morphological, and functional changes that evolve across stages of CNS pathology (Linnerbauer et al., 2020; Sofroniew, 2020; Escartin et al., 2021). Furthermore, astrocytes and microglia are the primary cellular mediators of the brain’s innate immune system, orchestrating the balance between immunity and neuroinflammation following insult, although increasing evidence also implicates the peripheral immune system (Ransohoff and Brown, 2012; De Rivero Vaccari and Keane, 2025; Duran et al., 2025; Tumpa et al., 2025).

Heavy alcohol consumption has been associated with increased neuroinflammatory responses in a number of preclinical and clinical studies (Crews and Vetreno, 2014; Cruz et al., 2023). Significant neuroimmune activation by alcohol has been shown in rodent models of excessive drinking and alcohol-dependence, such as the Chronic Intermittent Ethanol–Two-Bottle Choice (CIE-2BC) model and chronic alcohol exposures by intragastric gavage (Crews et al., 2013; Salem et al., 2024; Hashimoto et al., 2025a). While several studies have investigated the roles of microglia in alcohol-induced neuroinflammation (Fernandez-Lizarbe et al., 2009; Zhang et al., 2018; Lowe et al., 2020; Warden et al., 2020), the contribution of astrocytes in alcohol-related neuroimmune response remains poorly defined (Guizzetti et al., 2025). Earlier studies identified neuroimmune pathways impacted by CIE (Erickson et al., 2019) in astrocytes of the prefrontal cortex (PFC) that were not identified with chronic ethanol drinking (Erickson et al., 2018). Single-cell and single nuclei transcriptomic studies in the PFC have begun to elucidate interactions among astrocytes, microglia, oligodendrocytes, and neuronal populations in response to CIE-2BC (Salem et al., 2024). A recent study identified microglial and astrocytic neuroimmune and neuroinflammatory responses in human post-mortem dorsolateral PFC from individuals with Alcohol Use Disorder (AUD) (Warden et al., 2026). Although several studies from our group and others have examined the effects of chronic alcohol consumption and dependence on astrocyte gene expression and function in other brain regions, the impact of CIE-2BC on central amygdala (CeA) astrocytes remains unknown (Erickson et al., 2018, 2019; Hashimoto et al., 2025a; Nentwig et al., 2025; Hashimoto et al., 2026).

The CeA is a key brain region that integrates alcohol reward and stress responsivity, and plays a significant role in the development of alcohol dependence (Silberman et al., 2009; Roberto et al., 2010; Gilpin et al., 2015). Using rodent and non-human primates, we have demonstrated significant changes in cytokines and neuroimmune factors (e.g. IL-1b, IL-18, IL-6 and IL-10) that impact CeA function via modulation of inhibitory GABAergic transmission (Bajo et al., 2014, 2015, 2019, 2026; Harris et al., 2017; Roberto et al., 2017; Patel et al., 2019, 2021, 2022; Roberts et al., 2019; Cruz et al., 2023; Varodayan et al., 2023). In this study, by integrating multi-omic and morphometric approaches, we identified the molecular and morphological signature of CeA reactive astrocytes in ethanol dependence and identified astrocyte-specific mechanisms and molecular targets that can be developed into astrocyte-specific therapeutics for AUD.

## 2. Materials And Methods

### 2.1 Animals

Adult male and female B6;FVB-Tg(Aldh1l1-EGFP/Rpl10a)JD130Htz/J mice (astro-TRAP mice) (Doyle et al., 2008; Heiman et al., 2008) were purchased from The Jackson Laboratory (strain # 030247). Animal husbandry was as previously described (Hashimoto et al., 2025a). The Scripps Research Institute Institutional Animal Care and Use Committee approved all animal procedures following both ARRIVE, and US National Institutes of Health animal welfare guidelines.

### 2.2 Chronic intermittent ethanol – two bottle choice (CIE-2BC)

The mouse CIE-2BC model of alcohol dependence-induced escalation of drinking is widely accepted in the alcohol field to model the dependence-like alcohol consumption associated with AUD (Becker and Lopez, 2004; Chu et al., 2007; Finn et al., 2007; Warden et al., 2020; Patel et al., 2021; Borgonetti et al., 2023, 2025; Siddiqi et al., 2023; Salem et al., 2024). We carried out two independent cohorts of CIE-2BC studies in astro-TRAP mice; each cohort included 10-12 mice/sex exposed to CIE-2BC (Dep mice); 10-12 mice/sex exposed to 2BC (Non-Dep mice), and 6-12 mice/sex that were naïve to ethanol (Naïve mice). Mice were singly housed and acclimated to ethanol through 4 days (Mon-Fri) of 24hr access to 15% ethanol in addition to their normal food and water. The CIE-2BC procedure including blood ethanol concentration (BEC) determinations are previously described (Hashimoto et al., 2025a) and available in Supplementary Methods.

### 2.3 Translating ribosome affinity purification (TRAP) and RNA isolation

Frozen whole brains were sectioned (300µm) and bilateral brain punch microdissections were collected from the CeA using a 1.5 mm brain punch between Bregma -0.94 mm to -2.18 mm (Franklin and Paxinos, 2001). Tissue punches were placed into pre-chilled 1.7 mL microcentrifuge tubes and stored at -80° C until processing as previously described (Hashimoto et al., 2025a), with additional details in Supplementary Methods. Four sets of TRAP isolations (passes) were completed, each balanced by sex and treatment group (Dep, Non-Dep, Naïve).

### 2.4 RNA-Seq and analysis

The OHSU Integrated Genomics Laboratory (IGL) was used for library preparation and sequencing of RNA from TRAP and input samples as previously described (Hashimoto et al., 2025a). A total of 97 samples were sequenced (8 female Naïve, 8 male Naïve, 8 female Non-Dep, 7 male Non-Dep, 10 female Dep, 8 male Dep; in IP and input fractions) with one sample (input male Non-Dep sample) excluded prior to sequencing due to insufficient RNA yield. RNA-Seq processing and analysis was conducted as previously described (Hashimoto et al., 2025a), with additional details in Supplementary Methods. Differential expression/translation analysis for each dataset was performed using DESeq2 v1.26.0 (Love et al., 2014). The model used for the contrasts of “Dep vs Non-Dep”, “Dep vs Naïve”, and “Non-Dep vs Naïve” included Sex and TRAP pass as covariates. Additional comparisons were performed separately within each sex, using TRAP pass as a covariate. Significant regulation was determined by Benjamini-Hochberg (BH) multiple-comparison adjusted *p*-values < 0.05. To identify genes with higher expression in astrocytes (IP) versus the bulk-tissue, “IP vs input” comparisons were performed on the combined dataset within each treatment (Dep, Non-Dep, Naïve) using sex and TRAP pass as covariates. For IP vs Input comparisons, significant enrichment in the IP (astrocyte) fraction was determined by an adjusted *p*-value < 0.05 and a log_2_ fold-change > 1.

### 2.5 TRAP qRT-PCR

Confirmation of astrocyte specific mRNA enrichment by the TRAP procedure was carried out using qRT-PCR on a subset of the sequenced samples. A total of 16 input and IP samples (9 female, 7 male) from Non-Dep animals were used as previously described (Hashimoto et al., 2025b; Kawa et al., 2025). Input and IP RNA aliquots were analyzed using the Luna Universal One-Step RT-qPCR Kit (NEB) on the CFX96 (Bio-Rad) thermocycler with normalization to total RNA using the Quant-it RiboGreen kit (ThermoFisher). Primer sequences for *Il33*, *Aldh1l1*, *Tubb3, Itgam*, and *Mbp* were previously described (Talabot-Ayer et al., 2012; Hashimoto et al., 2025b).

### 2.6 Bioinformatic analysis

Ingenuity Pathway Analysis (IPA, Qiagen) was used for gene category enrichment analysis to determine enriched canonical pathways. Canonical pathway enrichment analysis produces p-values corresponding to the significance of the enrichment, enrichment ratio corresponding to the number of enriched genes in the analyzed dataset divided by the total number of genes in the pathway, and activation z-score which predicts activation or inhibition of pathways based on the direction of regulation of genes in the dataset.

### 2.7 Fluorescent in situ hybridization (RNAscope)

Fresh frozen brains were sectioned on a Leica CM3050S Cryostat and 16 µm sections were collected on Superfrost plus slides. RNAscope probes for *Aldh1l1* (Cat. # 405891-C2)*, Tubb3* (Cat. # 423391-C3), and *Stat1* (Cat. # 479611) or *Stat3* (Cat. # 425641) was run on brain sections according to the manufacturer’s recommendations excluding the protease step using the RNAscope Multiplex Fluorescent Reagent Kit v2 with TSA Vivid Dyes (ACD Bio, Cat. # 323270). Nuclei were counter stained with DAPI (Molecular Instruments; Los Angeles, CA, USA). Slides were imaged using a Leica DM500b microscope with a DFC365 FX camera and 6-8 images were taken of the CeA (bilaterally) using 40X objective lens from each brain section (1 section per animal per probe). Images were analyzed for co-expression of genes of interest (*Stat1* or *Stat3*) with cell-type specific markers. The total number of cells positive for astrocytic (*Aldh1l1*), neuronal (*Tubb3*), and gene of interest (*Stat1* or *Stat3*) markers were collected per image. The total number of dots for the gene of interest (*Stat1* or *Stat3*) were also counted for each *Aldh1l1* positive cell.

### 2.8 Proteomic & phosphoproteomic analyses

CeA were subjected to proteomic and phosphoproteomic analysis as previously described (Hardesty et al., 2022) and in Supplementary Methods. The mass spectrometry raw files were processed in a batch mode and searched together using DIA-NN (Demichev et al., 2020) (Version 1.8.1) against UniProt fasta of mouse proteome (2023-03-01-reviewed with contaminants, 21,949 entries) in default parameters with several modifications described in Supplementary Methods. The protein quantification was log_2_ transformed and statistical analysis was performed using linear regression modeling with the “limma” R package including sex as a covariate (Smyth, 2005), and *p*-values were adjusted by Benjamini-Hochberg (BH) procedure. For the proteomic analyses, we used a BH corrected *p-*value < 0.1 for significance thresholding. Due to the increased variability in phosphoproteomic signal and to capture more potential phosphorylation signals, for the phosphoproteomics we utilized an uncorrected *p*-value < 0.01. A total of 30 CeA samples were used for proteomics/phosphoproteomic analyses (n = 5, per sex, and treatment group) with both proteomic and phosphoproteomic analysis run on each sample.

### 2.9 Morphometric analysis of astrocytes

Whole brains of 9 female mice (n=3 Naïve, n=3 Non-Dep, and n=3 Dep) were rapidly removed, flash frozen in isopentane and stored at -80° C. Brains were thawed to room temperature and post-fixed in a 30% sucrose and 4% paraformaldehyde solution for 48hr. The CeA was sectioned at 60 μm (+2.2mm to -0.94mm from Bregma) (Paxinos and Watson, 2^nd^ Ed.) using a Leica SM 2000R sliding microtome and placed in phosphate buffered saline (PBS). Immunohistochemical amplification of endogenous EGFP was carried out using a free-floating section protocol using a 1:800 dilution rat anti-EGFP (Nacalai Cat#: 04404-84, RRID: AB_10013361) followed by a 1:500 dilution of Alexa Fluor donkey anti-Rat 488 (Thermo Fisher Scientific, Cat#: A-21208, RRID: AB_2535794) secondary antibody.

The CeA was imaged using a Zeiss LSM 780 confocal microscope (Zeiss, Oberkochen, Germany). Four bilateral images per animal were acquired with a 63X oil-immersion objective (NA 1.40). Z-stacks spanning 49–53 µm in depth were collected with a step size of 0.38 µm. Image stacks were imported into Neurolucida 360 (V2024.2.2; MBF Bioscience, Williston, VT) and the image background intensity was corrected using a rolling ball radius of 15. Approximately 6-9 reconstructions were completed per animal (1-4 astrocytic cellular reconstructions per image). Once reconstructions were completed, the data were exported to Neurolucida Explorer (V2022.2.1; MBF Bioscience, Williston, VT) to generate morphometric analyses and quantitative measurements of the astrocytic reconstructions. The sample size for astrocyte morphological assessments was a total of 19-20 cells per treatment group. A detailed description of immunohistochemistry and morphometric astrocyte reconstruction is available in Supplementary Methods.

### 2.10 Statistics

For the BEC data (obtained from Dep mice only), effects of sex, CIE week (i.e. the 5 different weeks of CIE), and sex by CIE week interaction were assessed by two-way repeated measure ANOVA. Measurements taken throughout the 2BC drinking (ethanol g/kg, body weight, ethanol mL, and water mL) were analyzed in the R package lme4 (Bates et al., 2015) using a linear mixed effects model to account for the multiple measurements within each week, with intercept-only models as previously described (Goeke et al., 2018). Effects of sex, ethanol vapor exposure (CIE), and sex by ethanol vapor exposure interactions were assessed by likelihood ratio tests with *p*-values generated by chi-squared test, where significance is determined as *p* < 0.05. Statistics for astrocyte reconstructions were performed using GraphPad Prism software (V10.5.0). Normality was assessed for each morphological measure within groups using the Shapiro–Wilk test. When normality assumptions were violated, group differences were assessed using Kruskal–Wallis tests with post-hoc Dunn’s multiple-comparison. Measures that approximated normal distributions were analyzed on the original scale using a one-way analysis of variance (ANOVA) with Tukey’s Multiple Comparisons test. Complexity measures were treated separately because two of the three groups significantly deviated from normality and variance increased with increasing complexity, data were log_10_-transformed prior to statistical analysis to stabilize variance. Moreover, analyses on the log scale allow group effects to be interpreted as proportional (fold-change) differences. Analysis where a 2-factor design was considered, (i.e., Sholl intersections or branch hierarchy and group) a two-way repeated measures (RM) ANOVA was performed with the Geisser-Greenhouse correction to accommodate missing values at distal radii or higher branch orders for some cells. Post-hoc Šídák’s multiple comparisons test was performed when an interaction was observed. Data were considered significant when p<0.05. Overlap between proteomics and RNA-Seq datasets were assessed using the R function *phyper* (R Core Team, 2025).

## 3. Results

### 3.1 CIE-2BC-induced escalation of drinking in Astro-TRAP mice

The CIE-2BC model of dependence induces escalation in drinking in mice exposed to ethanol vapor (Dep mice), but not in mice that voluntarily drink ethanol in the two-bottle-choice protocol (Non-Dep mice), making this a robust model of alcohol dependence where the escalation in ethanol drinking validates each experiment (Griffin et al., 2009). A schematic description of the CIE-2BC exposure used in these studies is shown in **Figure 1A**. **Figure 1B** shows the overall experimental design and the endpoints measured. Two independent cohorts of astro-TRAP mice were used for CIE-2BC studies. Each cohort included 10–12 mice per sex in the CIE-2BC group (Dep), 10–12 mice per sex in the 2BC group (Non-Dep), and 6–12 ethanol-naïve mice per sex (Naïve). The behavioral results of the first cohort of animals used in the bulk (input) RNA-Seq and in astro-TRAP-Seq analyses were previously described (Hashimoto et al., 2025a). Similarly, the second cohort of mice showed escalation in drinking after the 3^rd^ and 4^th^ CIE cycles (**Figure 1 C**), providing a strong behavioral rationale for the investigation of the effects of alcohol dependence on the CeA astrocyte translatome, proteome, and morphology. Each CIE cycle resulted in intoxicating blood alcohol levels (162-211mg/dl) (**Figure 1D**).

**Figure 1.**
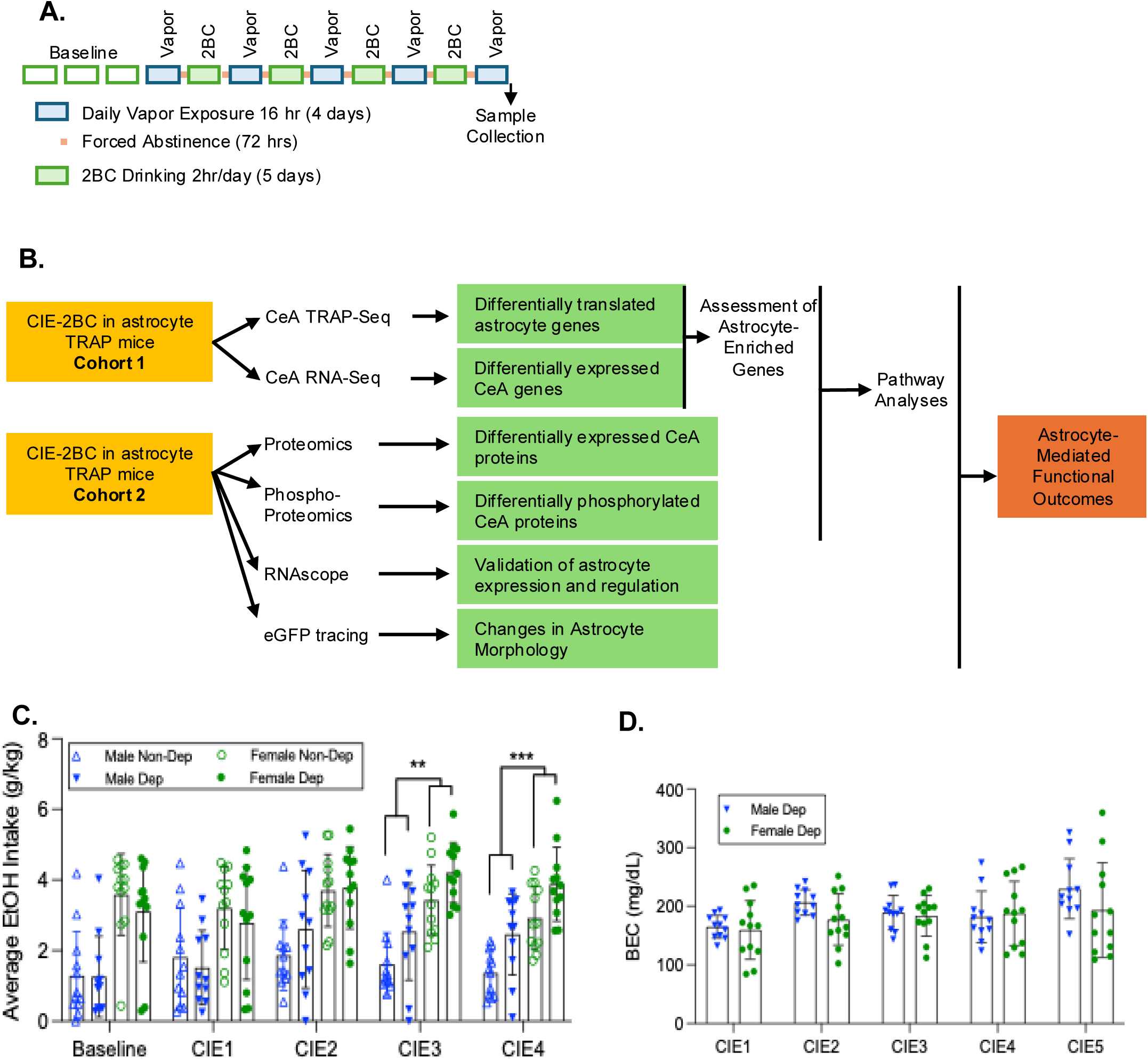
Experimental design and CIE-2BC behavioral outcome. **A.** Time-line of CIE-2BC procedure to induce ethanol dependence. The experimental design included astro-TRAP (Aldh1l1-EGFP/Rpl10a) mice exposed to the CIE-2BC procedure (Dependent, Dep), astro-TRAP mice that underwent 2BC voluntarily alcohol drinking (Non-Dependent, Non-Dep), and astro-TRAP ethanol-naïve mice (Naïve). **B.** Schematic of the experimental design and endpoints measured. **C.** Average ethanol intake at baseline and after CIE weeks 1 to 4 in Dep mice compared to Non-Dep mice showing escalation of drinking following CIE week 3 (CIE3) and CIE week 4 (CIE4). **D.** Blood ethanol concentrations (BECs) following each CIE in Dep mice showing consistent BECs across the 5 CIE exposures. **C** and **D** are from the second cohort of CIE-2BC-exposed mice (11-12 animals/sex/treatment); escalation in drinking and BECs from the first cohort of mice has been described in (Hashimoto et al., 2025a).

### 3.2 Astrocyte translatome analysis by TRAP-Seq

The astrocyte translatome was evaluated in astro-TRAP mice expressing an EGFP tag on ribosomal protein RPL10a driven by the astrocyte-specific *Aldh1l1* promoter that allows for the pull-down of astrocyte-specific translating RNA (Doyle et al., 2008; Heiman et al., 2008). The selective pull-down of astrocyte RNA after the TRAP protocol in comparison to the bulk CeA RNA (input) was verified by qPCR with TRAP samples showing enrichment in the astrocyte marker *Aldh1l1* and depletion in neuronal (*Tubb3*), microglial (*Itgam*), and oligodendrocyte (*Mbp*) markers (**Supplemental Figure 1A**), consistent with our previous results using this technique (Hashimoto et al., 2025b, 2025a).

CeA astrocyte-specific translating mRNA changes and bulk tissue transcriptional changes were assessed in Dep, Non-Dep, and Naïve astro-TRAP-mice by TRAP-Seq and RNA-Seq, respectively. The average RNA Integrity Number (RIN) for all samples was 7.6 ± 0.197 (**Supplemental Table 1**).

Hierarchical clustering of all the sequenced samples shows clear separation of input (bulk CeA mRNA) and IP (astro-TRAP, astrocyte translating RNA) samples (**Supplemental Figure 1B**). Similarly, principal component analysis (PCA) shows input and IP samples separating along the x-axis (principal component 1) in **Supplemental Figure 1C**, with 81% of the sample variance. PCA of the IP samples only did not reveal any clear separation based on treatment group or sex but did show some separation based on the TRAP experiment batch (**Supplemental Figures 1D-F**). To assess whether ethanol treatments altered the expression of the transgene driven by the *Aldh1l1* promoter, we determined the reads aligning to EGFP and found no significant alteration of EGFP based on sex, treatment, or sex by treatment interactions in the input or IP samples (data not shown).

### 3.3 Ethanol dependence induces changes consistent with neuroimmune activation and alterations in antioxidants/oxidative stress pathways while non-dependent ethanol drinking alters cytoskeleton plasticity pathways in astrocyte translating RNA

The number of significantly regulated genes based on BH adjusted (adj) *p*-values in the comparisons Dep vs Non-Dep, Non-Dep vs Naïve, and Dep vs. Naïve from the astro-TRAP-Seq and bulk CeA (input) RNA-Seq datasets are summarized in **Table 1**. In all comparisons, there were more DT genes in the astro-TRAP fractions than DE genes in the input fractions, indicating that the cell-type specific TRAP-Seq approach can unveil differences in astrocyte-specific mechanisms that are obscured in bulk RNA-Seq studies. In this paper, we report male- and female-combined analyses; the complete list of differentially translated (DT; TRAP-Seq) and differentially expressed (DE; RNA-Seq) genes in analyses of combined and separate sexes are reported in **Supplemental Table 2**. We sequenced both the input RNA and the astrocyte translating RNA to be able to identify genes that are predominantly astrocytic. We calculated astrocyte-enriched genes defined as genes significantly more expressed (adj p<0.05) and with a log_2_ fold-change (FC) > 1 in the astrocyte TRAP fraction compared to the input fraction (**Supplemental Table 3**). The number of astrocyte-enriched genes differentially regulated by treatments is shown in the right column of **Table 1**. **Supplemental Figure 2** shows the Venn diagrams of the overlap of the astro-TRAP DT (**A**) and of the input DE (**B**) genes across treatments.

**Table 1.** Number of differentially translated and differentially expressed genes. Astro-TRAP: astrocyte translating RNA from the CeA. Input: bulk CeA tissue. Dep: Dependent; Non-Dep: Non-Dependent. Regulated genes: adjusted p-value < 0.05. Regulated astrocyte-enriched genes are regulated genes that are more expressed in astro-TRAP fraction than in the input fraction with a >1 log_2_ fold-change and adj p<0.05.

| Fraction | Comparison | Regulated Genes | Upregulated Genes | Down-regulated Genes | Regulated Astrocyte-Enriched Genes |
| --- | --- | --- | --- | --- | --- |
| Astro-TRAP | Dep vs Non-Dep | 384 | 249 | 135 | 188 |
| Astro-TRAP | Non-Dep vs Naïve | 296 | 175 | 121 | 150 |
| Astro-TRAP | Dep vs Naïve | 938 | 509 | 429 | 328 |
| Input | Dep vs Non-Dep | 68 | 60 | 8 | 30 |
| Input | Non-Dep vs Naïve | 141 | 68 | 73 | 26 |
| Input | Dep vs Naïve | 544 | 309 | 235 | 97 |

Ingenuity Pathway Analysis (IPA) was used to identify canonical pathways with significant overrepresentation in the astro-TRAP datasets. The CIE-2BC protocol results in increased voluntary drinking (referred to as alcohol dependence); to infer astrocyte-mediated processes involved in alcohol dependence, the comparison Dep vs Non-Dep is of particular interest. **Figure 2A** shows the top 30 IPA categories overrepresented in the Dep vs Non-Dep comparison, all of which relate to immune functions (highlighted in yellow) and most showing a positive activation z-score, indicating activation of inflammatory pathways. **Figure 2B** shows genes (and the cellular location of their gene products) regulated in the Dep vs Non-Dep comparison that are involved in the “Inflammatory response” function based on the IPA Interpret feature “Diseases and Functions”. Upregulated genes are in red/pink, down-regulated genes are in blue/light blue, and genes shown in bold are enriched in astrocytes.

**Figure 2.**
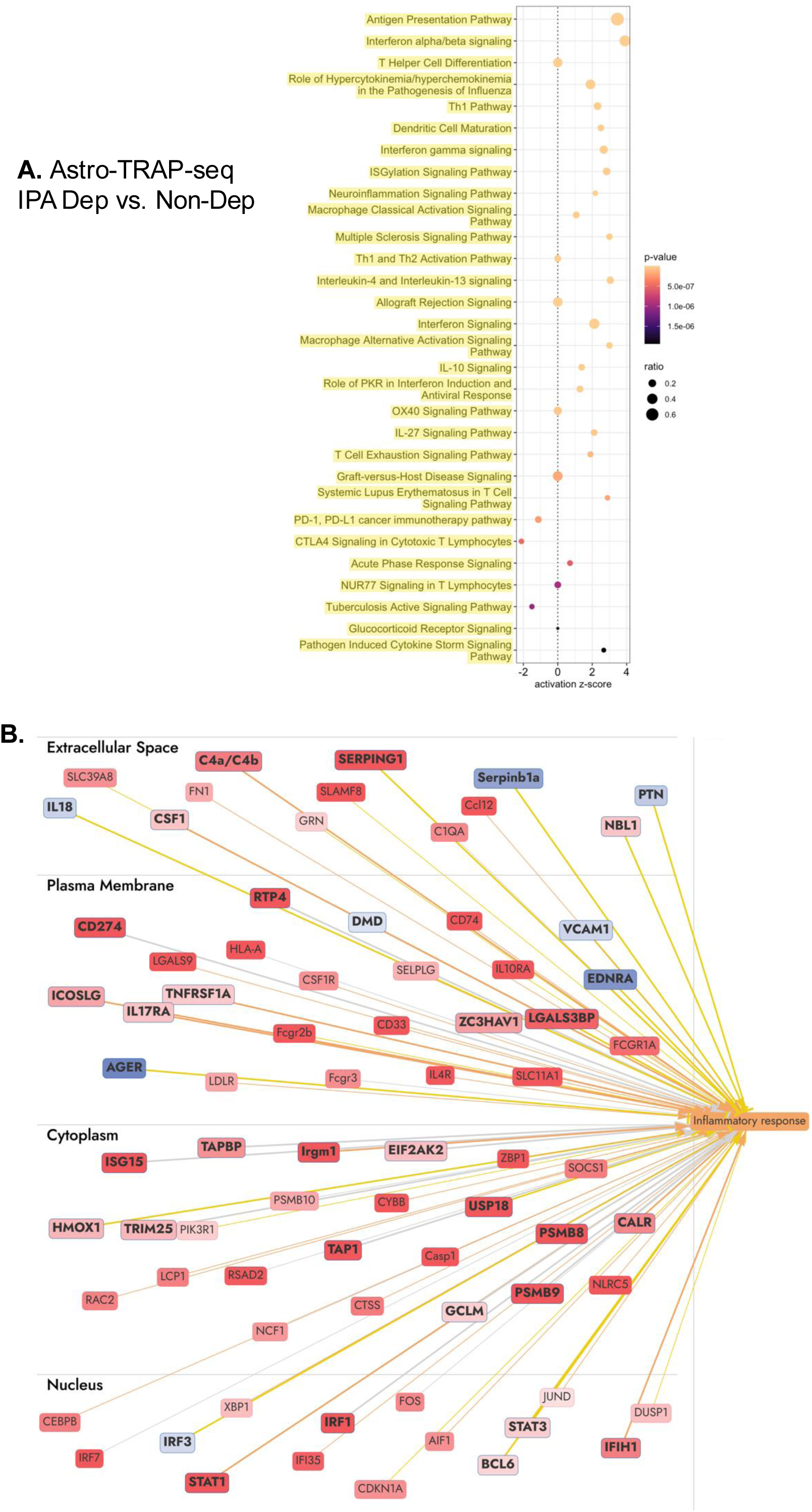
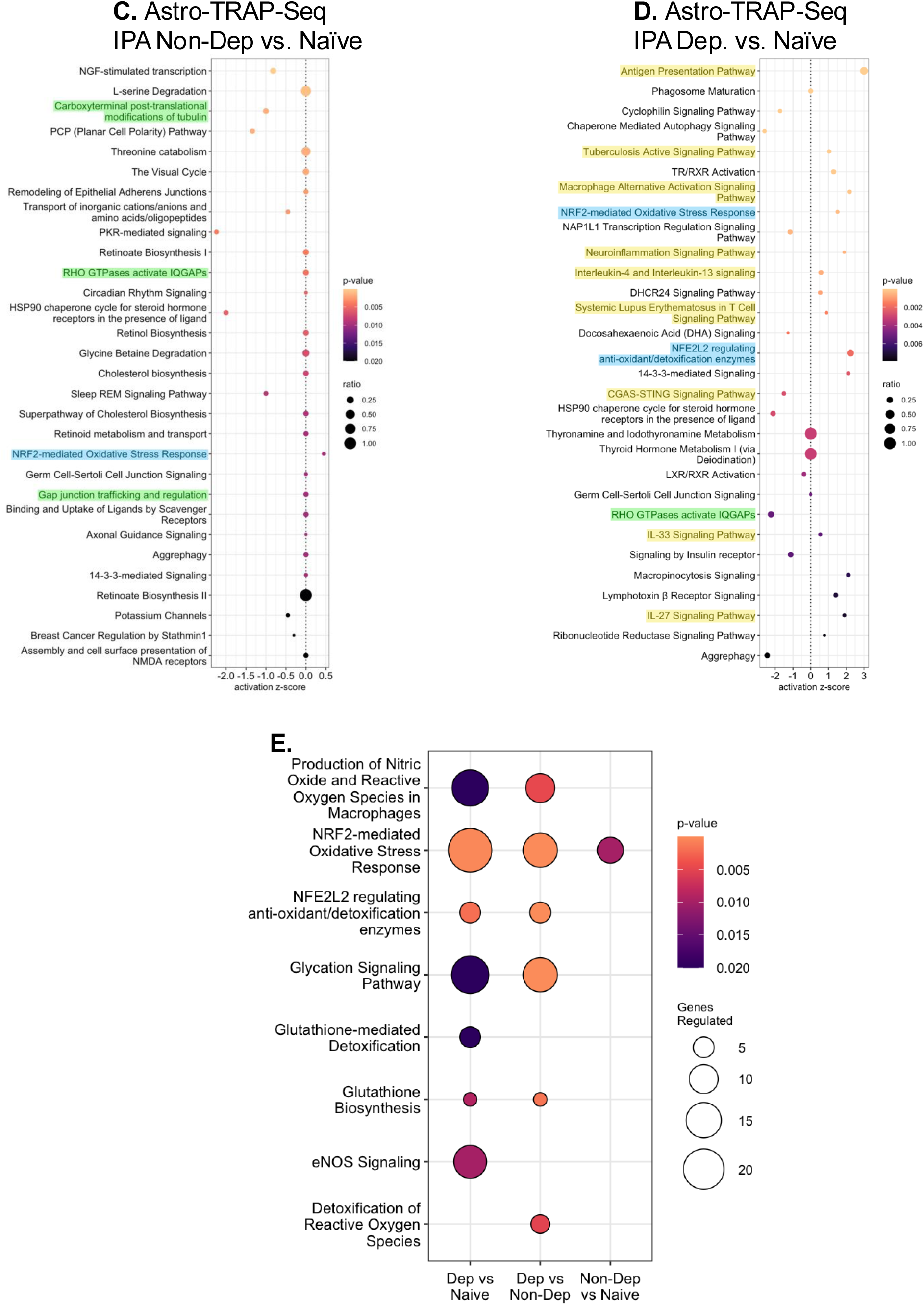
IPA overrepresented categories in astro-TRAP-Seq datasets. **A.** The top 30 overrepresented IPA canonical pathways in the Dep vs. Non-Dep comparison are all related to neuroimmune/neuroinflammation functions (highlighted in yellow) with most showing a positive activation z-score, indicating neuroimmune activation. **B.** “Inflammatory Response” IPA function includes 76 genes regulated in the Dep vs Non-Dep comparison and the subcellular location of the proteins encoded by these genes. The “Inflammatory Response” function was predicted to be significantly activated in Dep mice compared to Non-Dep mice by the IPA Interpret feature “Diseases and Functions”. Genes in pink-to-red are upregulated (low-to-high); genes in blue are down-regulated; the orange box and orange arrows predict activation; blue arrows predict inhibition; genes in bold are enriched in astrocytes. **C.** The top 30 overrepresented IPA canonical pathways identified in astro-TRAP-Seq results from the Non-Dep vs. Naïve comparison include pathways involved in cytoskeleton remodeling (highlighted in green). **D.** The top 30 overrepresented IPA canonical pathways identified in astro-TRAP-Seq results from the Dep vs. Naïve comparison show activation of several neuroimmune/neuroinflammation and antioxidant/oxidative stress pathway. In panels **A**, **C**, and **D**: the x-axis shows the activation z-score which is a prediction of overall pathway activation or inhibition based on known pathway relationships and the regulated genes in the pathway, the size of each circle represents the proportion (ratio) of DT genes to the total pathway size, and the color of each circle corresponds to the pathway overrepresentation p-value, with brighter colors corresponding to lower p-values. Pathways highlighted in yellow are neuroimmune/neuroinflammatory related pathways, blue are oxidative stress related pathways, and green are cytoskeleton related pathways. **E**. Comparison of oxidative stress related pathways with enrichment p-values < 0.05 are shown in the Dep vs Naïve, Dep. vs Non-Dep, and Non-Dep vs Naïve comparisons. For each comparison shown on the y-axis, the circle size corresponds to the number of genes differentially translated in astrocytes and the circle color corresponds to the p-value of the overrepresented category.

Voluntary ethanol drinking also elicited a high number of changes in astrocyte translating RNA levels compared to ethanol Naïve mice (**Table 1**). The top 30 IPA categories overrepresented in the Non-Dep vs Naïve comparison are shown in **Figure 2C**. Significantly overrepresented IPA pathways did not show neuroimmune activation by ethanol drinking, instead, some pathways identified are involved in the remodeling of the cytoskeleton (highlighted in green), suggesting ethanol drinking may induce morphological changes. It should be noted that, in contrast to pathways in the Dep vs Non-Dep comparison in which immune pathways are mostly activated (as indicated by positive activation z-scores), most of the regulated pathways in the Non-Dep vs Naïve comparison are homeostatic, as indicated by the activation z-scores around 0.

The Dep vs Naïve comparison yielded the highest number of regulated genes in the astro-TRAP-Seq dataset (**Table 1**), consistent with these being the most divergent experimental conditions and with previous observations in the NAc (Hashimoto et al., 2025a). The top 30 IPA categories overrepresented in the Dep vs Naïve comparison shown in **Figure 2D** include immune activation, cytoskeleton remodeling, and oxidative stress/antioxidant system pathways.

Astrocytes play a key role in protecting the brain from oxidative stress and are major contributors to antioxidant systems. However, upon transition to a reactive state, astrocytes downregulate antioxidant functions and begin producing ROS. **Figure 2E** shows the oxidative stress/antioxidant system-related IPA pathways overrepresented in the three comparisons and reveals that alcohol dependence strongly alters pathways involved in the modulation of oxidative stress in astrocytes. The full list of significant canonical pathways in the three comparisons and their activation z-score is available in **Supplemental Table 4**. Together, these results indicate that in the progression from non-dependent alcohol consumption to alcohol dependence, astrocytes undergo neuroimmune activation and activation of pathways involved in the regulation of redox status.

### 3.4 Ethanol dependence induces neuroimmune activation and alterations in antioxidants/oxidative stress pathways while ethanol drinking alters cytoskeleton plasticity pathways in proteomics studies

As a complementary approach to our RNA-Seq analyses, we ran LC-MS/MS proteomics on CeA samples from Dep, Non-Dep, and Naïve astro-TRAP mice to investigate changes in protein expression. Since the proteomics analysis is not astrocyte-specific, we leveraged astro-TRAP-Seq vs input RNA-Seq data to infer cell-type specificity. We identified regulated proteins that are enriched in astrocytes and also applied an additional filtering step to exclude proteins whose genes show significantly lower gene expression in astrocytes than in the bulk CeA (i.e. proteins that likely are regulated in non-astrocyte cell types). **Table 2** summarizes the number of regulated proteins in each of these groups. The lists of significantly regulated proteins (total, astrocyte-enriched, and regulated proteins minus proteins that show less expression in astrocytes compared to the bulk CeA) are shown in **Supplemental Table 5**.

**Table 2.** Number of differentially expressed proteins.

| Comparison | # regulated proteins |  |  | # regulated proteins enriched in astrocytes |  |  | # regulated proteins minus proteins low expressed in astrocytes |  |  |
| --- | --- | --- | --- | --- | --- | --- | --- | --- | --- |
|  | Total | Up | Down | Total | Up | Down | Total | Up | Down |
| Dep vs Non-Dep | 289 | 220 | 69 | 61 | 41 | 20 | 128 | 88 | 40 |
| Non-Dep vs Naïve | 266 | 68 | 198 | 41 | 28 | 13 | 78 | 61 | 17 |
| Dep vs. Naïve | 145 | 95 | 50 | 41 | 35 | 6 | 84 | 40 | 44 |

**Figure 3A** & **B** shows the top 30 IPA canonical pathways overrepresented in the Dep vs Non-Dep comparison in the complete proteomic dataset (**A**) and after removing proteins less expressed in astrocytes (**B**). The complete lists of significantly overrepresented IPA canonical pathways in the three comparisons of the two datasets are shown as **Supplemental Tables 6 and 7**. Both analyses were characterized by the presence of several overrepresented pathways related to immune responses (in yellow) and to oxidative stress/antioxidant systems (in blue) with several of these categories having a positive activation z-score and none with a negative activation z-score, in agreement with the astro-TRAP-Seq analyses (**Figure 2**). In addition, several categories related to cholesterol biosynthesis, trafficking, and lipid homeostasis were identified (highlighted in orange), suggesting that alcohol dependence alters the production and trafficking of cholesterol and other lipids.

**Figure 3.**
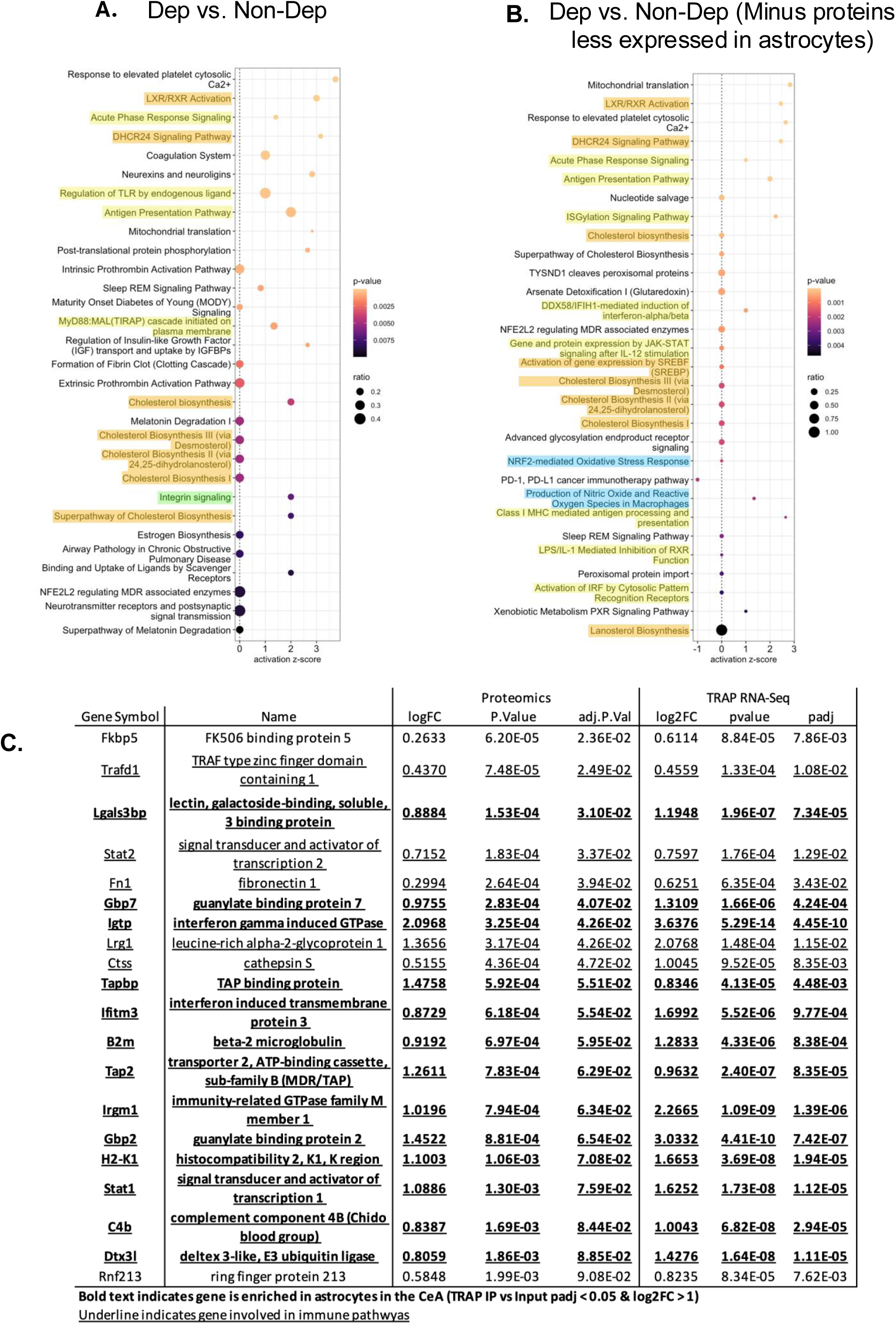
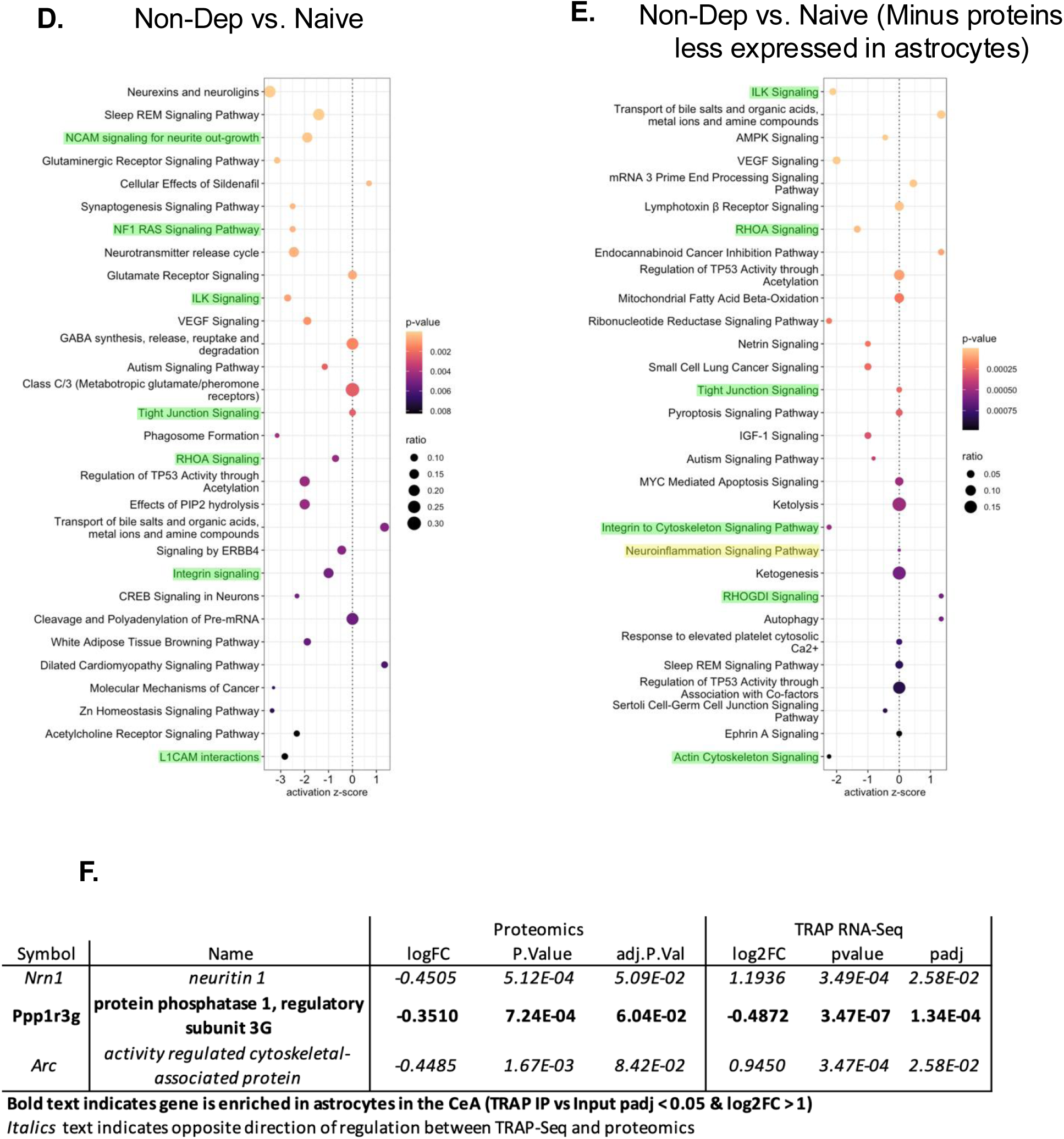
Pathway analysis of CeA proteomics. **A.** Top 30 IPA canonical pathways from the Dep vs. Non-Dep comparison. **B.** Top 30 IPA canonical pathways from the Dep vs. Non-Dep comparison with regulated proteins minus regulated proteins whose genes are low expressed in astrocytes compared to the bulk tissue. **C.** List of overlapping changes in expression of genes (by astro-TRAP-Seq) and proteins (by proteomic analysis) from the Dep vs. Non-Dep comparison. **D.** Top 30 IPA canonical pathways from the Non-Dep vs. Naïve comparison. **E.** Top 30 IPA canonical pathways from the Non-Dep vs. Naïve comparison with regulated proteins minus regulated proteins whose genes are low expressed in astrocytes compared to the bulk tissue. In panels **A-B** and **D-E**, the x-axis shows the activation z-score which is a prediction of overall pathway activation or inhibition based on known pathway relationships and the regulated proteins in the pathway, the size of each circle represents the proportion (ratio) of significant proteins to the total pathway size, and the color of each circle corresponds to the enrichment p-value, with brighter colors corresponding to lower p-values. Pathway names related to cholesterol are highlighted in orange, immune responses are highlighted in yellow, oxidative stress/antioxidant are highlighted in blue, and cytoskeleton are highlighted in green. **F.** List of overlapping changes in expression of genes (by astro-TRAP-Seq) and proteins (by proteomic analysis) from the Non-Dep vs Naïve comparison.

We next compared the overlap in mRNA (astro-TRAP-Seq) and protein (proteomics) regulation and identified 20 genes in both datasets, all were regulated in the same direction; this represents a significant overlap in hypergeometric testing (q=20, k=289, m=384, n=14,823; *p*=4.7x10^-5^). Thirteen of these genes/proteins are enriched in astrocytes and 18 of them are involved in immune responses (**Figure 3C**). The concordant upregulation in astro-TRAP-Seq and proteomics of the same genes and proteins related to immune functions indicates a persistent upregulation of neuroimmune functions in astrocytes.

The top 30 overrepresented categories in the Non-Dep vs. Naïve comparison using the entire proteomic dataset and after removing proteins whose genes are significantly less expressed in astrocytes are shown in **Figure 3D** and **E**, respectively. In agreement with the astro-TRAP-Seq analyses, these comparisons identify cytoskeleton-related pathways as significantly altered by alcohol drinking; however, overlap in differential expression of individual genes and proteins was minimal (**Figure 3F**).

The top 30 categories in the complete proteome dataset and after removing proteins less expressed in astrocytes in the Dep vs Naïve comparison are shown in **Supplemental Figure 3A & B**.

### 3.5 Convergent evidence indicates persistent upregulation of the astrocyte-specific complement gene/protein *C4b*/C4 by alcohol dependence

*C4b* and its encoded protein C4, a member of the complement system also involved in synaptic pruning (Zhou et al., 2022; Zheng et al., 2025), are consistently upregulated by alcohol dependence (**Figure 4A, B, C**). *C4b* is primarily expressed by astrocytes in the CeA, with some astrocytes expressing very high levels of *C4b* as shown in a representative RNAscope image (**Figure 4D**). The high level of expression of *C4b* in astrocytes relative to other cell types in the CeA is also evident when comparing expression of *C4b* in the astro-TRAP-Seq and the input RNA-Seq fractions (**Figure 4E**). Additional complement genes, *C1qa*, *C1qb*, and *C1qc*, and *Serping1*, encoding for the C1 inhibitor C1-INH protein, were upregulated in the astro-TRAP-Seq Dep vs Non-Dep comparison; these genes are less expressed in astrocytes than in the input (**Figure 4E**) and were not regulated in the input RNA-Seq or in the proteomic analyses (**Figure 4F**).

**Figure 4.**
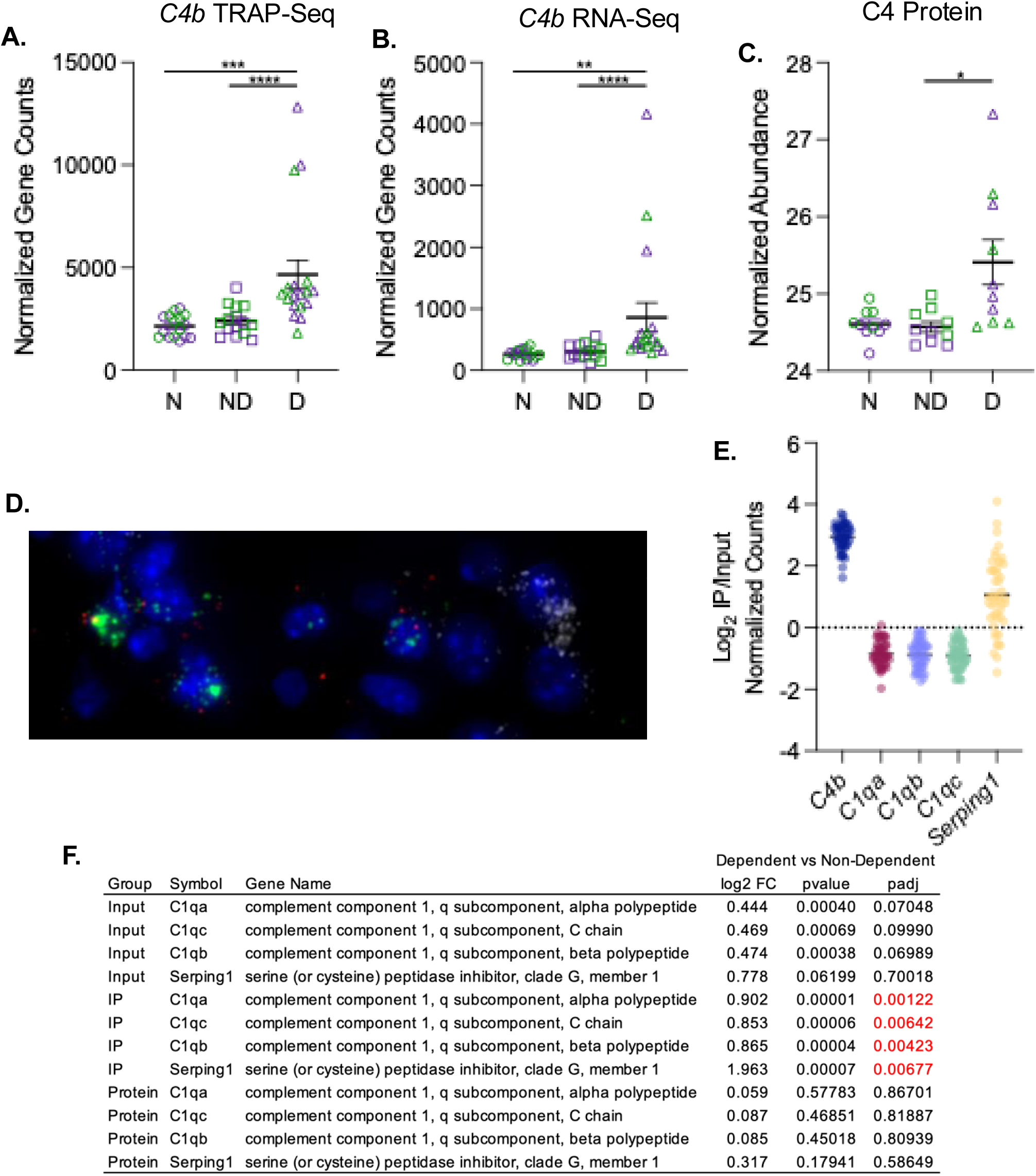
Effect of alcohol dependence on complement genes and proteins. Dependent (D) animals had higher levels of C4b than Non-Dep (ND) and Naïve (N) animals in the CeA in the astrocyte (IP) fraction (**A**) and bulk CeA RNA (Input) fraction (**B**). Protein analysis of C4 protein levels showed significant elevation in Dep vs Non-Dep animals (**C**). **D**. Representative RNAscope image shows C4b (green dots) mRNA localized to astrocytes expressing Aldh1l1 (red dots), and little C4b localization in neurons expressing Tubb3 (white dots). Nuclei are counter-stained with DAPI (blue). **E.** Relative expression of complement related genes in CeA astrocytes vs bulk CeA RNA shows astrocyte enrichment of C4b (3.04 log_2_ fold-change, adj p-value = 3.8x10^-43^) and decreased astrocyte expression of C1qa (-0.96 log_2_ fold-change, adj p-value = 2.3x10^-9^), C1qb (-0.88 log_2_ fold-change, adj p-value = 8.5x10^-8^), and C1qc (-0.96 log_2_ fold-change, adj p-value = 1.8x10^-8^). Serping1 shows a trend for increased expression in astrocytes (0.79 log_2_ fold-change, adj p-value = 0.098). **F.** Table of complement related mRNA and protein expression. The significance thresholds for RNA data (Input and IP Groups) are adj p < 0.05 while for proteins the threshold is adj p < 0.1.

### 3.6 Convergent evidence indicates astrocyte STAT activation by alcohol dependence

The Signal Transducer and Activator of Transcription (STAT)1, 2, 3 proteins, encoded by *Stat1*, *Stat2*, and *Stat3* genes, are transcription factors that are regulated by neuroimmune signaling, including interferon signaling pathways (Dong et al., 2022). We found that these three *Stat* genes are upregulated in astro-TRAP-Seq analyses in the Dep vs Non-Dep comparison (**Figure 5A**). *Stat1* is also upregulated in the input RNA-Seq (**Figure 5B**), and STAT1 and STAT2 proteins are upregulated in proteomic analysis (**Figure 5C**). The increased expression of *Stat1* in astrocytes in the Dep vs Non-Dep comparison was confirmed by RNAscope (**Figure 5D, E**). *Stat1* and *Stat3* genes are enriched in astrocytes, while *Stat2* is expressed at similar levels in astrocytes and in the bulk tissue (**Figure 5F**).

**Figure 5.**
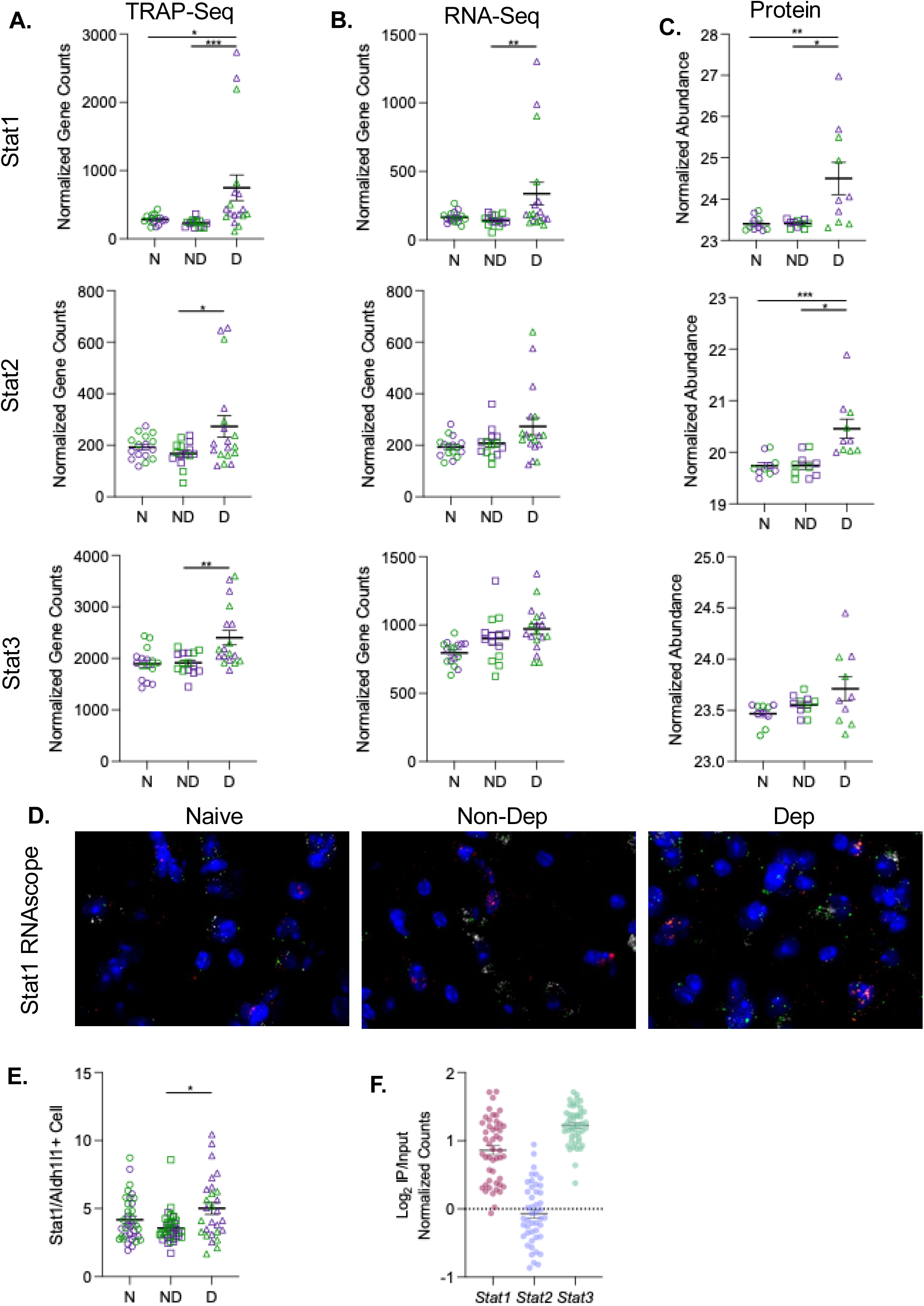
Ethanol dependence alters Stat1, Stat2, and Stat3 expression. **A.** *Stat1*, *Stat2*, and *Stat3* translation was upregulated by alcohol dependence in CeA astrocytes (TRAP-Seq). **B**. *Stat1*, but not *Stat2* and *Stat3* expression was upregulated in the bulk CeA (RNA-Seq) of dependent mice. **C.** STAT1 and STAT2 protein was increased in the CeA of Dep mice. **D.** Representative RNAscope images in CeA from Naïve, Non-Dep, and Dep animals. Stat1 mRNA is green, astrocyte marker Aldh1l1 is red, neuronal marker Tubb3 is white, and nuclei, counter-stained with DAPI, is blue. **E.** Analysis of RNAscope images show elevated number of Stat1 dots in Aldh1l1 positive astrocyte cells (*p = 0.05). **F.** Relative expression of Stat genes in CeA astrocytes vs bulk CeA RNA shows elevated astrocyte expression of Stat1 (0.78 log_2_ fold-change, adj p-value = 0.005) and Stat3 (1.25 log_2_ fold-change, adj p-value = 2.8x10^-58^), and no difference in Stat2 expression in the astrocyte (IP) fraction compared to the bulk-tissue (input) expression (0.002 log_2_ fold-change, adj p-value = 0.99). Plotted are all samples log_2_ (IP normalized counts/Input normalized counts). D: dependent; ND: Non-Dep; N: Naïve mice.

### 3.7 Proteins involved in astrocyte physiological functions are downregulated by alcohol dependence

Evidence presented so far indicates that astrocytes from Dep mice upregulate neuroimmune and oxidative stress/antioxidant pathways, consistent with the transition toward a reactive astrocyte phenotype. This transition is also characterized by downregulation of physiological astrocyte functions (Escartin et al., 2021). We have found evidence of significant downregulation of astrocyte-specific genes and proteins that are known to play important physiological roles. The astrocyte-enriched gene *Hapln1* encoding for the key perineuronal net component hyaluronan and proteoglycan link protein 1 is downregulated in the astro-TRAP-Seq Dep vs Non-Dep comparison (log_2_ fold-change: -0.429; adj *p*=0.0058). The astrocyte-enriched gene *Ptn* encoding for the protein pleiotrophin, is downregulated in astro-TRAP-Seq in the Dep vs Non-Dep comparison (log_2_ fold-change: -0.397; adj *p*=0.0078). Pleiotrophin plays an important role in dendrite and synaptic structure (Brandebura et al., 2025), its deficiency is associated with depressive-like behavior (Chi et al., 2025) and its overexpression reduces adolescent alcohol consumption (Galán-Llario et al., 2024). The astrocyte protein connexin 43 (CX43), a key component of astrocyte gap junctions allowing the rapid transfer of ions, gliotransmitters, and second messengers between neighboring astrocytes, encoded by the gene *Gja1* is downregulated in proteomics studies (log_2_ fold-change: -0.383; adj *p*=0.042). The astrocyte K^+^ channel Kir4.1 (encoded by the gene *Kcnj10*), important for K^+^ reuptake after action potential, displays significant downregulation of S345 phosphorylation in Dep vs Non-Dep mice (log_2_ fold-change: - 1.6; *p*=0.0002). All these changes are consistent with inhibition of astrocyte physiological functions by alcohol dependence.

### 3.8 Ethanol drinking and ethanol dependence induce changes in the phosphorylation of proteins involved in cytoskeleton remodeling

The activity and function of proteins involved in intracellular processes, requiring rapid activation and inactivation, are often modulated by reversible phosphorylation. We conducted phosphorylation site-level comparisons to characterize the differential phosphorylation regulation in CeA samples from Naïve, Non-Dep, and Dep mice. **Table 3** shows the number of phosphorylation sites and **Supplemental Table 8** shows the complete list of phosphorylation sites significantly altered in the three comparisons. To identify specific treatment-induced effects on protein phosphorylation and rule out protein phosphorylation changes that are secondary to changes in protein expression, we removed phosphorylated protein sites that were regulated in the same direction as the observed changes in protein abundance prior to canonical pathway analyses (**Supplemental Table 8**). To determine the canonical pathways impacted by the CIE-2BC treatments on protein phosphorylation, IPA was used. The top 30 pathways are shown in **Supplemental Figure 4** and the full list of significantly overrepresented pathways are available in **Supplemental Table 9**.

**Table 3.** Number of phosphorylation sites altered by ethanol drinking or ethanol dependence.

| Comparison | Up-Regulated | Down-Regulated | Total Regulated |
| --- | --- | --- | --- |
| Dep vs Non-Dep | 94 | 77 | 171 |
| Non-Dep vs Naïve | 248 | 60 | 308 |
| Dep vs. Naïve | 225 | 43 | 268 |

\*Counts are based on sites of phosphorylation; proteins can be modified at multiple sites

Canonical pathways identified in phosphoproteomic datasets from the Non-Dep vs Naïve, Dep vs Naïve, and, to a lesser extent, Dep vs Non-Dep comparisons included pathways involved in cytoskeleton remodeling suggesting that alcohol drinking induces cytoskeletal remodeling that may be further enhanced during the development of alcohol dependence. Interestingly, the strong neuroimmune signature observed in the astro-TRAP-Seq and proteomic analyses in dependent mice, was absent in the phosphoproteomic analyses indicating that neuroimmune molecules are primarily regulated through gene and protein expression rather than protein phosphorylation. **Figure 6A** shows the significant IPA canonical pathways related to cytoskeletal remodeling, including several categories related to RHO-GTPases, involved in cytoskeletal remodeling and morphological changes in astrocytes (Domingos et al., 2023), across the three comparisons.

**Figure 6.**
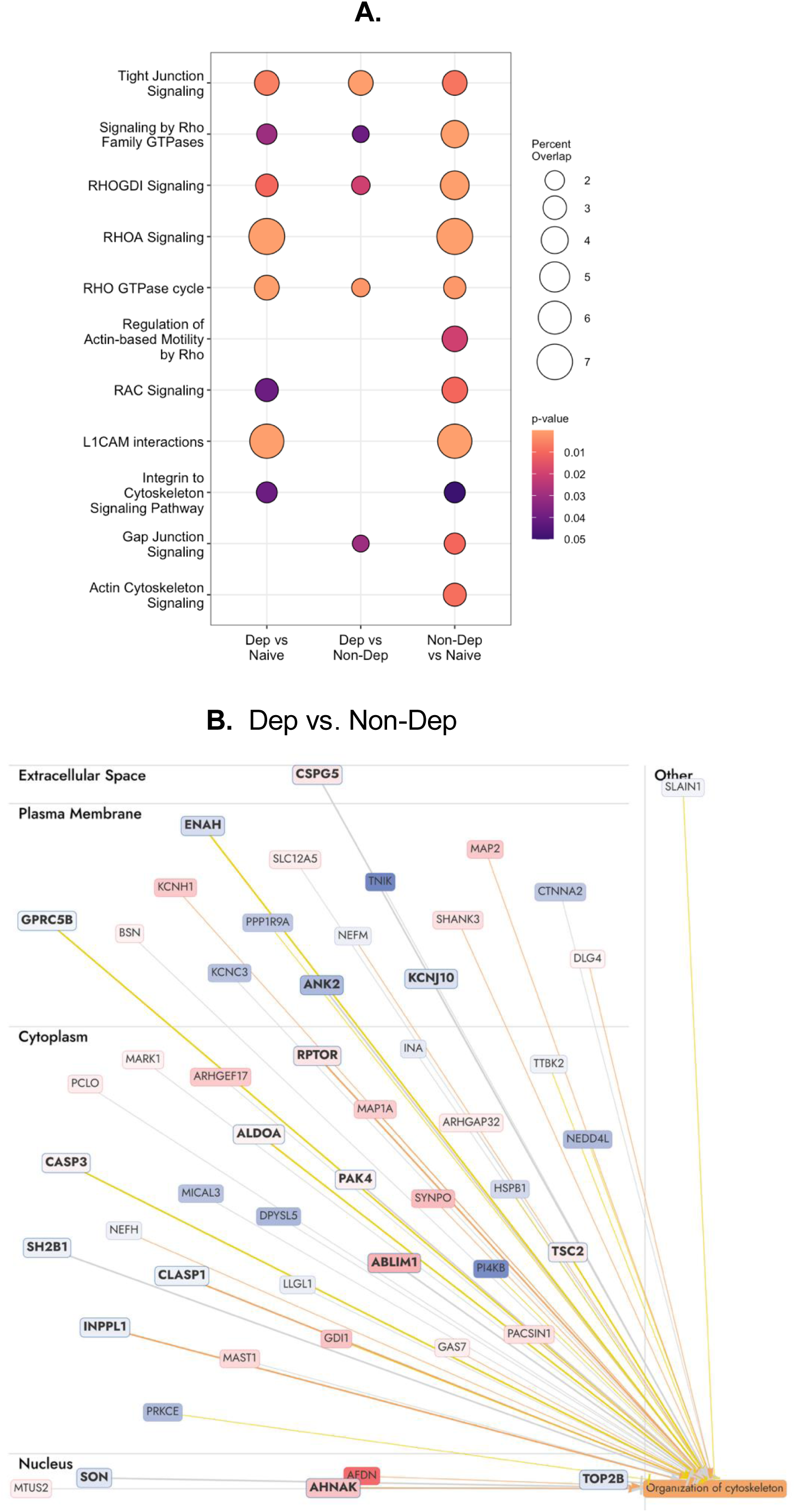
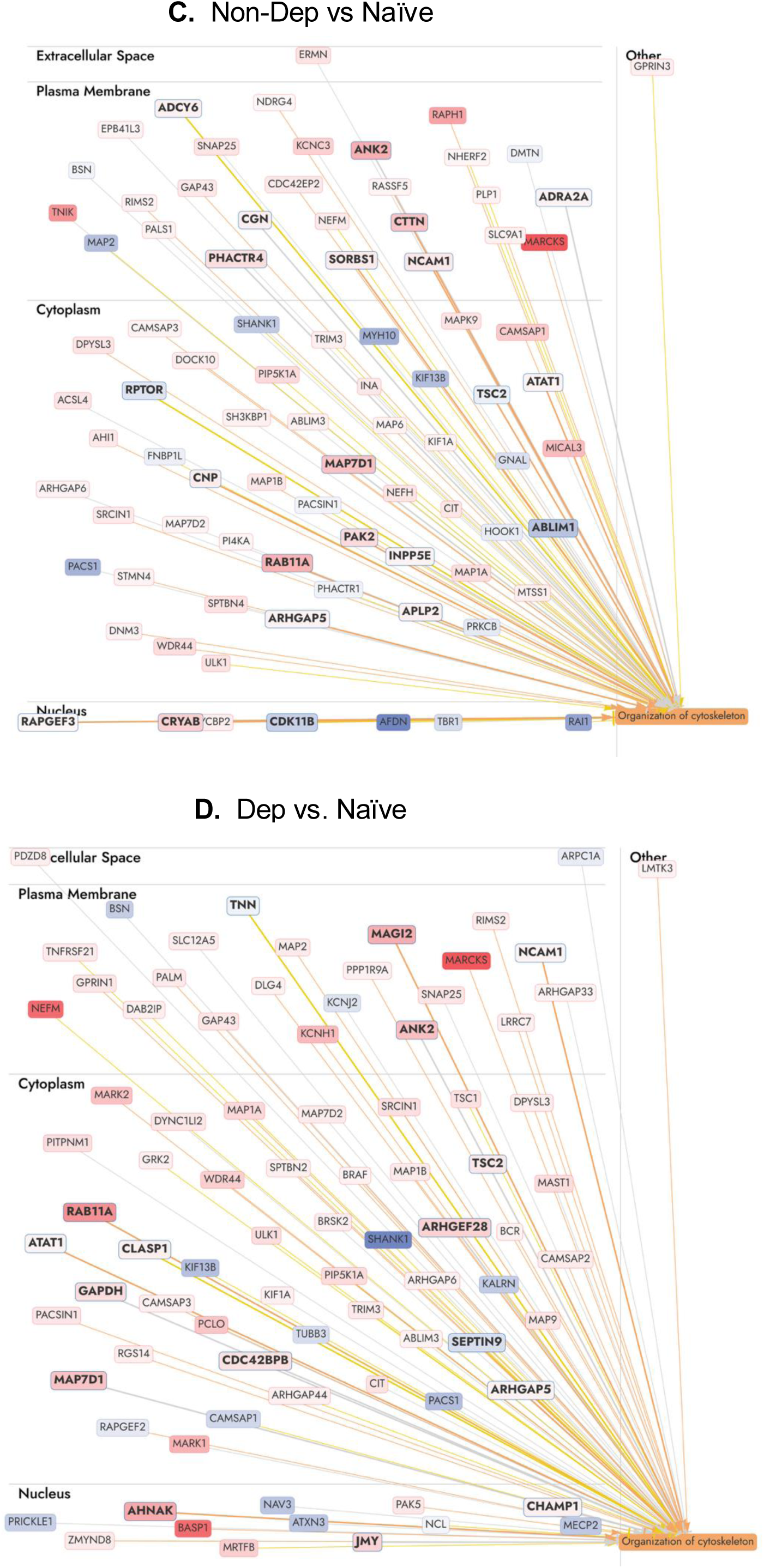
A. Cytoskeleton related pathways altered by ethanol in phosphoproteomics analysis. Differential phosphorylation results were analyzed in IPA for canonical pathway overrepresentation. IPA phosphoproteomics analysis utilizes protein level phosphorylation patterns to identify pathways with more phosphorylated proteins than expected by chance. Pathways with enrichment p-values < 0.05 were selected and pathways relating to cytoskeleton remodeling are shown in the Dep vs Naïve, Dep vs Non-Dep, and Non-Dep vs Naïve comparisons (**A**). For each pathway shown on the y-axis, the circle size corresponds to the percent overlap of the differentially phosphorylated proteins in each of the three comparisons shown on the x-axis compared to the total pathway size. The color of the circle corresponds to the enrichment p-value with brighter colors being more significant. In cases where the pathway did not meet the enrichment significance threshold (p < 0.05) no circle is plotted. **B-D**. Shown is the “Organization of Cytoskeleton” function that includes 51 proteins regulated in the Dep vs Non-Dep (**B**), 86 proteins regulated in the Non-Dep vs Naïve (**C**), and 85 proteins regulated in the Dep vs Naïve (**D**) comparisons. Proteins in pink-to-red are upregulated (low-to-high); proteins in blue are down-regulated; the orange box and orange arrows predict activation; blue arrows predict inhibition; proteins in bold are encoded by genes that are either significantly more expressed in astrocytes or are expressed in astrocytes and the bulk tissue at similar levels; the other proteins are encoded by genes significantly less expressed in astrocytes than in the bulk tissue.

To identify a broad range of proteins involved in cytoskeletal remodeling with altered phosphorylation, we employed the IPA feature “Diseases and Functions” that identified “Organization of cytoskeleton” as a significantly regulated function in all three comparisons and included 51 proteins in the Dep vs Non-Dep comparison (**Figure 6B**), 86 proteins in the Non-Dep vs Naïve comparison (**Figure 6C**), and 85 proteins in the Dep vs Naïve comparison (**Figure 6D**). Because the phosphoproteomic analyses are not cell type specific, and to gain insights on whether cytoskeleton remodeling occurs in astrocytes, we categorized proteins that are encoded by genes enriched in astrocytes, significantly more expressed in astrocytes, equally expressed in astrocytes as in the bulk CeA, and significantly less expressed in astrocytes using the astro-TRAP-Seq and input RNA-Seq data set as described above. **Figure 6B-D** shows the proteins in the “Organization of the cytoskeleton” function; proteins in bold text are either enriched in astrocytes or expressed at similar levels in astrocytes and the bulk tissue; these are proteins that are more likely to play a role in cytoskeleton remodeling in astrocytes. **Supplemental Table 10** shows all the altered phosphorylation sites in the proteins from the “Organization of cytoskeleton” IPA function.

### 3.9 Alcohol drinking and alcohol dependence alter CeA astrocyte morphology

Astro-TRAP-Seq, proteomic, and phosphoproteomic data suggest that alcohol drinking alters cytoskeletal remodeling pathways in astrocytes, whereas alcohol dependence primarily affects these pathways at the phosphoproteomic level. Cytoskeleton remodeling can lead to morphological changes in astrocytes which are often associated with astrocyte reactivity (Baldwin et al., 2023). Thus, using immunohistochemistry, confocal imaging, and morphometric analyses, we quantified the length of the main astrocyte branches, the number of branch bifurcations and endings, the overall cellular complexity, and the spatial territory (area and volume) occupied by CeA astrocytes from female Dep, Non-Dep, and Naïve astro-TRAP mice.

Astro-TRAP mice allow for accurate and quantitative measures of astrocyte morphology by leveraging the EGFP tag on astrocyte ribosomes for visualization of comparative morphology (Viana et al., 2023). **Figure 7A** shows EGFP-labeled CeA astrocytes from Naïve astro-TRAP mice. The number of main astrocyte processes did not differ among groups (data not shown). However, there was a significant difference among groups in the total process length and the number of nodes and endings as revealed by a non-parametric Kruskal-Wallis analysis (*p*<0.0001, length, nodes, endings). Dunn’s multiple comparisons showed increased process length and increased number of nodes and endings in both Non-Dep and Dep groups compared to Naïve mice, but no difference between Non-Dep and Dep (**Figure 7B, C, D**). One-way ANOVA revealed an effect of treatment on overall astrocytic complexity (F (2, 56) = 58.60, *p*<0.0001; **Figure 7E**).

**Figure 7.**
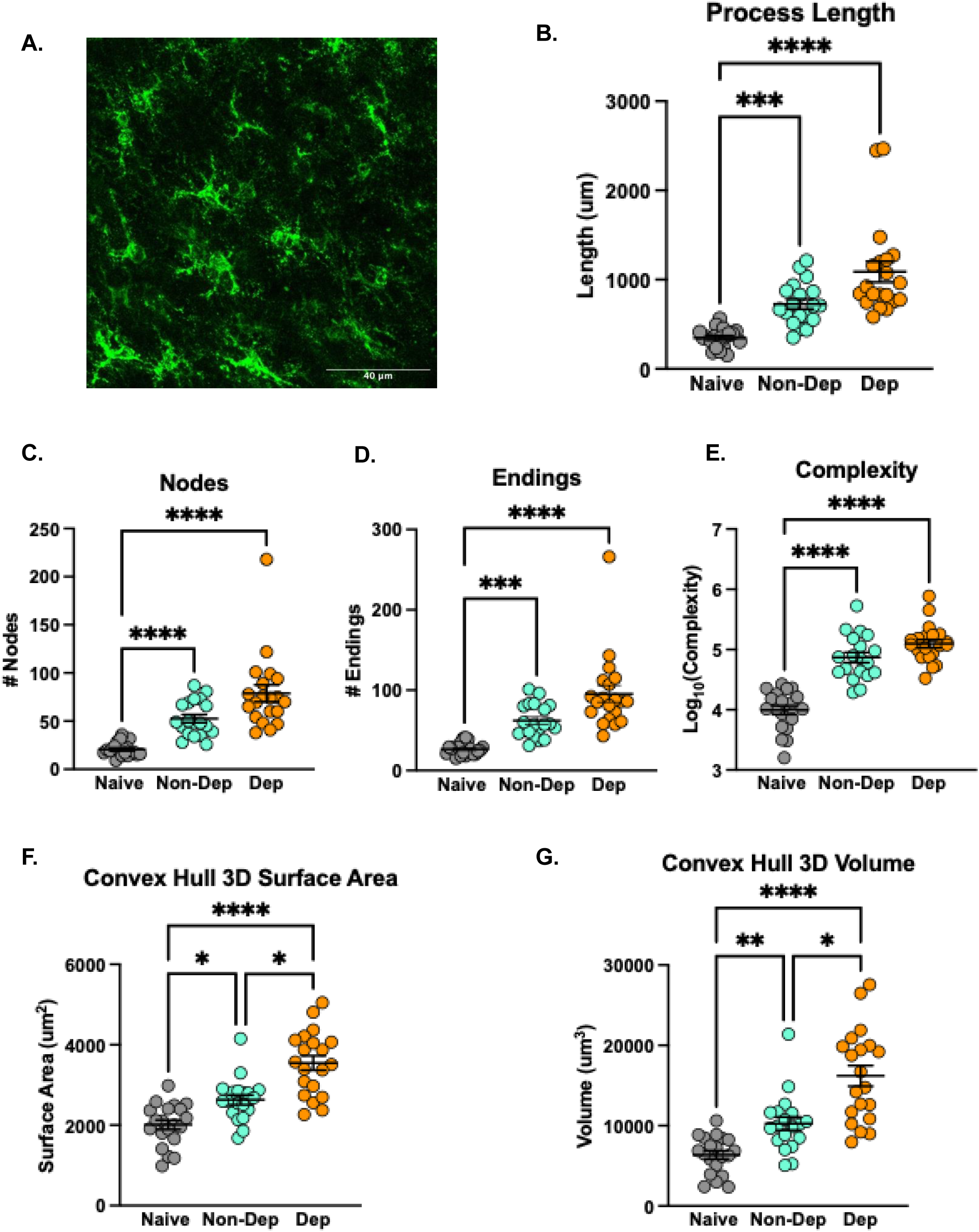
Effects of alcohol exposure and dependence on CeA astrocyte morphology. (**A**) Representative image of EGFP-amplified Aldh1l1-EGFP expression in Naïve CeA astrocytes. Structural comparison of CeA astrocytes (**B**) process length, (**C**) nodes, and (**D**) endings across Naïve, Non-Dep and Dep groups. (**E**) Analysis of astrocyte complexity across groups (One-Way ANOVA). (**F**) Comparison of astrocytic territory surface area (convex hull 3D). (**G**) Comparison of astrocytic territory volume (convex hull 3D). N=19-20 astrocytes (3 mice/group). Data are presented as mean ± SEM. Group differences were analyzed using Kruskal-Wallis test followed by Dunn’s multiple comparisons across groups unless otherwise stated. **\**p*** < 0.05, **\*\**p*** < 0.01, **\*\*\**p*** < 0.001, **\*\*\*\**p*** < 0.0001.

Tukey’s post-hoc analysis showed increased complexity in both Non-Dep and Dep groups compared to Naïve mice, while no differences were observed between Non-Dep and Dep groups. Astrocyte territory coverage using 3D convex hull analysis revealed an effect of treatment on territory surface area and volume (**Figure 7F, G**, both *p*<0.0001) and Dunn’s post-hoc analysis showed increased territory surface area and volume in Non-Dep and Dep groups compared to Naïve and in the Dep compared to the Non-Dep groups.

To examine astrocyte branching complexity, we assessed the highest branch order across groups using Kruskal-Wallis analysis (**Figure 8A**, *p<*0.0001). Post-hoc Dunn’s multiple comparisons revealed that both Non-Dep and Dep groups achieved higher branch orders compared to Naïve mice. Next, branch level changes were analyzed using Two-way RM ANOVA. The number of branches across branch orders showed a significant interaction effect (**Figure 8B**, Branch Order x Treatment F (7.419, 207.7) = 11.13, *p*<0.0001). Tukey’s post-hoc analysis showed that compared to Naïve, the Non-Dep group had increased number of branches from orders 3 to 14, while the Dep group had increased number of branches from orders 1 to 12. The Dep mice also exhibit increased number of branches from orders 3 to 7 compared to the Non-Dep mice. Branch length across orders revealed an interaction (**Figure 8C**, Two- way RM ANOVA, Branch Order x Treatment F (10.14, 283.9) = 6.866, *p*<0.0001), and post-hoc Tukey’s multiple comparisons showed increased branch length in the Non-Dep group across orders 6 to 14 and across orders 3 to 14 for the Dep group relative to Naïve. Further, there was increased branch length across orders 3, 6, and 7 in the Dep vs Non-Dep groups.

**Figure 8.**
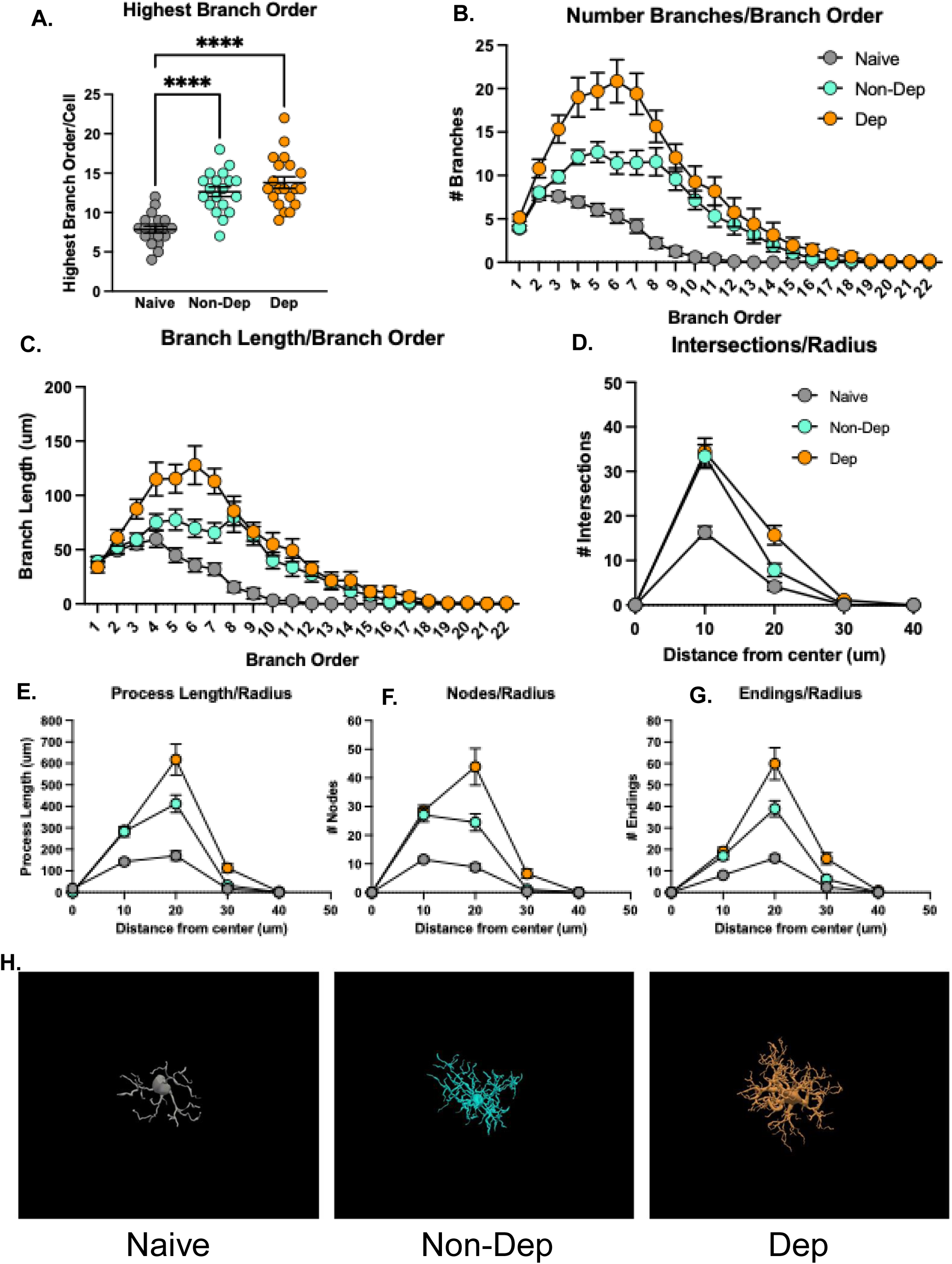
CeA astrocyte branch order and Sholl analyses in Naïve, Non-Dep, and Dep groups. Using the centrifugal ordering schematic, the branching hierarchy was assigned by the number of segments traversed from the root to describe the complexity of astrocyte arborization. (**A**) Maximum branch order achieved by treatment group (Kruskal-Wallis test) **\*\*\*\****p* < 0.0001. (**B-C**) Quantitative comparison of arbor complexity at each level of centrifugal ordering. (**B**) number of branches/branch order (Interaction p<0.0001, Effect of Order p<0.0001, Effect of Treatment p<0.0001; Order 1: Naïve vs Dep p=0.0462, Order 2: Naïve vs Dep p=0.0379, Order 3: Naïve vs. Non-Dep p=0.0404, Naïve vs. Dep p=0.0379, Non-Dep vs. Dep p=0.0113; Order 4: Naïve vs. Non-Dep p<0.0001, Naïve vs Dep p=0.0001, Non-Dep vs. Dep p=0.0223; Order 5: Naïve vs. Non-Dep p=0.0001, Naïve vs Dep p<0.0001, Non-Dep vs. Dep p=0.0186; Order 6: Naïve vs. Non-Dep p=0.0008, Naïve vs Dep p<0.0001, Non-Dep vs. Dep p=0.0058; Order 7: Naïve vs. Non-Dep p=0.0003, Naïve vs Dep p<0.0001, Non-Dep vs. Dep p=0.0191; Order 8: Naïve vs. Non-Dep p<0.0001, Naïve vs Dep p<0.0001, Non-Dep vs. Dep p=0.2321; Order 9: Naïve vs. Non-Dep p<0.0001, Naïve vs Dep p<0.0001; Order 10: Naïve vs. Non-Dep p<0.0001, Naïve vs Dep p=0.0004; Order 11: Naïve vs. Non-Dep p=0.0029, Naïve vs Dep p=0.0004; Order 12: Naïve vs. Non-Dep p=0.0047, Naïve vs Dep p=0.0077; Order 13: Naïve vs Dep p=0.016, Order 14: Naïve vs Dep p=0.0389). (**C**) branch length/branch order (Interaction p<0.0001, Effect of Order p<0.0001, Effect of Treatment p<0.0001; Order 3: Naïve vs. Dep p=0.0118, Non-Dep vs. Dep p=0.0338; Order 4: Naïve vs Dep p=0.0092; Order 5: Naïve vs. Non-Dep p=0.0214, Naïve vs Dep p=0.0001; Order 6: Naïve vs. Non-Dep p=0.00611, Naïve vs Dep p=0.00012, Non-Dep vs. Dep p=0.0164; Order 7: Naïve vs. Non-Dep p=0.0108, Naïve vs Dep p<0.0001, Non-Dep vs. Dep p=0.0079; Order 8: Naïve vs. Non-Dep p=0.0007, Naïve vs Dep p=0.0001; Order 9: Naïve vs. Non-Dep p<0.0001, Naïve vs Dep p<0.0001; Order 10: Naïve vs. Non-Dep p=0.0004, Naïve vs Dep p=0.0003; Order 11: Naïve vs. Non-Dep p=0.0054, Naïve vs Dep p=0.0009; Order 12: Naïve vs. Non-Dep p=0.0036, Naïve vs Dep p=0.0004; Order 13: Naïve vs. Non-Dep p=0.0173, Naïve vs Dep p=0.0335; Order 14: Naïve vs. Non-Dep p=0.0469, Naïve vs Dep p=0.0388). Sholl analysis was used to assess (**D**) number of intersections/radius (Interaction p<0.0001, Effect of Radius p<0.0001, Effect of Treatment p<0.0001; 10µm: Naïve vs Non-Dep p<0.0001, Naïve vs Dep p<0.0001; 20µm: Naïve vs. Dep p=0.0001, Non-Dep vs. Dep p=0.0147; 30µm: Naïve vs. Dep p=0.0051, Non-Dep vs. Dep p=0.0152), (**E**) process length/radius (Interaction p<0.0001, Effect of Radius p<0.0001, Effect of Treatment p<0.0001; 10µm: Naïve vs Non-Dep p<0.0001, Naïve vs Dep p<0.0001; 20µm: Naïve vs. Non-Dep p<0.0001, Naïve vs. Dep p<0.0001, Non-Dep vs Dep p=0.0484; 30µm: Naïve vs. Dep p=0.0002, Non-Dep vs. Dep p=0.0017), (**F**) number of nodes/radius (Interaction p<0.0001, Effect of Radius p<0.0001, Effect of Treatment p<0.0001; 10µm: Naïve vs Non-Dep p<0.0001, Naïve vs Dep p<0.0001; 20µm: Naïve vs. Non-Dep p=0.0002, Naïve vs. Dep p<0.0001, Non-Dep vs Dep p=0.0284; 30µm: Naïve vs. Dep p=0.0028, Non-Dep vs. Dep p=0.0081), and (**G**) number of endings/radius (Interaction p<0.0001, Effect of Radius p<0.0001, Effect of Treatment p<0.0001; 10µm: Naïve vs Non-Dep p=0.0005, Naïve vs Dep p<0.0001; 20µm: Naïve vs. Non-Dep p<0.0001, Naïve vs. Dep p<0.0001, Non-Dep vs Dep p=0.0468; 30µm: Naïve vs. Non-Dep p=0.0177, Naïve vs. Dep p=0.0003, Non-Dep vs. Dep p=0.0081; 40µm: Naïve vs. Dep p=0.0127, Non-Dep vs. Dep p=0.0069). (**H**) Representative 3D renderings of astrocytes from Naïve, Non-Dep and Dep animals. N=19-20 astrocytes (3 mice/group). Data are presented as mean ± SEM. Group differences were analyzed using Two-way RM ANOVA followed by Tukey’s post-hoc analysis unless otherwise stated.

We also performed Sholl analysis to quantify changes in process complexity and spatial organization across different radii. Two-way RM ANOVA revealed an interaction for process intersections by radius (**Figure 8D**, Radius x Treatment F (3.095, 86.65) = 10.89, *p*<0.0001). Post-hoc Tukey’s multiple comparisons at 10 µm showed increased radii intersections for Non-Dep and Dep groups compared to the Naïve group, while at 20 and 30µm, the Dep group showed increased intersections compared to the Non-Dep and Naïve groups. Two-way RM ANOVA also revealed a radius x treatment interaction for process length (**Figure 8E**, F (2.750, 77.01) = 15.59, *p*<0.0001), number of nodes (**Figure 8F**, F (2.878, 80.59) = 14.58, *p*<0.0001), and number of process endings (**Figure 8G**, Radius x Treatment F (2.830, 79.23) = 14.99, *p*<0.0001). Post-hoc analysis showed increased process length at 10 and 20 µm for the Non-Dep compared to Naïve; the Dep group showed increased process length at 10, 20, and 30 µm compared to Naïve, and increased process length at 20 and 30 µm relative to the Non-Dep group. Post-hoc analysis also showed increased number of nodes per radii at 10 and 20 µm for the Non-Dep compared to Naïve; the Dep group showed increased nodes at 10, 20, and 30 µm compared to Naïve and at 20 and 30 µm compared to Non-Dep group. Tukey’s analysis showed increased process endings at 10, 20, and 30 µm for the Non-Dep group compared to Naïve; the Dep group showed increased process endings at 10, 20, 30, and 40 µm compared to Naïve and increased process endings at 20, 30, and 40 µm compared to Non-Dep group. **Figure 8H** shows representative 3D renderings of astrocytes from Naïve, Non-Dep and Dep animals. Statistical analyses for **Figures 7** and **8** are reported in **Supplemental Table 11**.

Together, these findings show that alcohol drinking results in structural remodeling in the CeA astrocytes with increased size and complexity. In addition, 3D convex hull metrics, Scholl, and branch order analyses revealed further significant differences between the Dep and Non-Dep groups, supporting the observation that alcohol dependence increases the area and volume of astrocytes already increased by alcohol drinking.

## 4. Discussion

In this study, we investigated the molecular and morphological responses of astrocytes to alcohol dependence using the established CIE-2BC paradigm, which results in escalation of drinking (Becker and Lopez, 2004; Warden et al., 2020; Patel et al., 2021; Borgonetti et al., 2023, 2025; Siddiqi et al., 2023; Salem et al., 2024). The study was carried out in astro-TRAP transgenic mice and included three experimental groups: mice naïve to ethanol, non-dependent mice that voluntarily consumed ethanol, and ethanol dependent mice exposed to ethanol vapor. Using astrocyte-specific translatomic, transcriptomic, proteomic, and phosphoproteomic analyses in combination with morphological assessment, we identified key features of astrocyte reactivity in the CeA of ethanol-dependent mice, indicating neuroimmune activation, oxidative stress- related changes, downregulation of physiological astrocyte functions, and cytoskeletal remodeling leading to morphological changes. Together, these findings demonstrate that alcohol dependence is associated with a shift toward a reactive astrocyte phenotype in the CeA and identify reactive astrocyte molecular and morphological signature specific to alcohol dependence.

Combining astro-TRAP-Seq and proteomics analyses, we found that the molecular signature of astrocytes from dependent mice is characterized by a preponderance of overrepresented pathways that predict activation of neuroimmune/inflammatory responses (**Figures 2**, **3; Supplemental Tables 2-7**), consistent with a transition from physiological astrocytes to reactive astrocytes (Lee et al., 2023). Reactive astrocytes are key components of CNS innate immunity and are highly heterogeneous, with molecular profiles that vary across brain regions, disease states, and stages of disease. While astrocyte reactivity is an evolutionary-conserved, adaptive response to brain insults, reactive astrocytes can become maladaptive and contribute to brain damage (Sofroniew, 2020). Pathway analyses of the dependent vs non-dependent comparisons revealed activation of neuroimmune and oxidative stress/antioxidant system functions in both proteomic and astro-TRAP-Seq analyses, confirming a persistent neuroimmune activation; we also observed statistically significant overlap between altered gene and protein expression. Most of the regulated genes and proteins, however, are not in common between the two datasets (**Figures 2**, **3**), which may be due to timing of protein synthesis and turnover and the dynamics of mRNA synthesis and degradation.

Our proteomic analyses indicate that cholesterol homeostasis is disrupted in dependent mice (**Figure 3 A, B**). While oligodendrocytes are the major producers of cholesterol in the brain, astrocytes also produce large amounts of cholesterol and astrocyte-derived cholesterol provides cholesterol to neurons for axon growth and synaptogenesis (Mauch et al., 2001; Hayashi et al., 2004; Valenza et al., 2015). Astrocytes also produce and release lipoproteins that contain cholesterol and ApoE (Vance and Hayashi, 2010; Pfrieger and Ungerer, 2011). Lipoproteins are trafficked to neurons via cholesterol transporters whose expression is regulated by LXR/RXR heterodimers, and we previously reported increased cholesterol trafficking in astrocytes exposed to ethanol *in vitro* (Guizzetti et al., 2007; Chen et al., 2013). *ApoE*, *Abca1*, *Lcat*, *Npc1*, *Npc2*, *Dhcr7*, genes involved in cholesterol biosynthesis and trafficking, are enriched in astrocytes (**Supplemental Table 3**), emphasizing the role of astrocytes in cholesterol biosynthesis and delivery to other cell types, most notably, to neurons.

The “complement system” pathway is significantly activated in the astro-TRAP analysis of dependent vs non-dependent mice, with *C1qa*, *C1qb*, *C1qc*, *C4b*, and *Serping1* all up-regulated. Upregulation of C1q proteins can lead to the activation of the complement cascade, which has been shown to occur in the brain (Tenner and Petrisko, 2025). However, the upregulation of *Serping1*, which encodes the C1 inhibitor C1-INH, in astrocytes suggests an inhibition of the complement cascade activation (**Figure 4**). In the brain, different complement components can be produced by different cell types and individual complement components in the brain influence cellular processes in the absence of the activation of the whole complement cascade (Tenner and Petrisko, 2025). Of note, *C1qa*, *C1qb*, and *C1qc* are typically thought of as being primarily expressed by microglia, but a recent single nuclei RNA-Seq analysis in the dorsal striatum detected all three transcripts in astrocyte nuclei with *C1qc* showing significant down-regulation 72 h after CIE exposure (Wildermuth et al., 2025). Based on our results, we hypothesize that components of the complement system upregulated in dependent mice have functions that are independent from the complement cascade activation. In particular, *C4b*/C4 expression, which is up-regulated in astro-TRAP-Seq, input RNA-Seq, and proteomic datasets (**Figure 4 A, B, C, F**) has been associated with neuroinflammation, phagocytosis, and neurodegenerative diseases (Goetzl et al., 2018; Winston et al., 2019; Dejanovic et al., 2022; Zhou et al., 2022; Nóbrega et al., 2025; Zou et al., 2025). In agreement with what has been reported by others, the *C4b* gene is predominantly expressed by astrocytes, with some astrocytes expressing very high levels of *C4b* mRNA (**Figure 4D, E**). The highly consistent and convergent findings of *C4b*/C4 upregulation in alcohol dependence highlights a potential role of *C4b*/C4 in the development of alcohol dependence and needs to be further explored.

Consistent with our findings in the nucleus accumbens of dependent mice (Hashimoto et al., 2025a), we identified dysregulation of Stat genes and proteins in the CeA of dependent mice. *Stat1*, *Stat2*, and *Stat3* are upregulated in the dependent group in the Astro-TRAP data set, STAT1 and STAT2 proteins were upregulated in the proteomic analysis, and *Stat1* and *Stat3* are enriched in astrocytes (**Figure 5**). STAT proteins are important players in neuroimmune responses where they can exert both pro- and anti-inflammatory effects and are down-stream effectors of cytokines and interferons (Hoffman et al., 2026). Several cytokine- and interferon-related pathways are overrepresented in the astro-TRAP and proteomic IPA analyses (**Figures 2**, **3**; **Supplemental Tables 4, 6, 7**) indicating that astrocyte *Stat1*(and possibly *Stat2* and *Stat3*) may be central to cytokine- and interferon-induced neuroimmune activation in dependent mice.

We also identified antioxidant-related pathways in the astrocyte translatome of dependent mice (**Figure 2; Supplemental Table 4**), that were also confirmed by proteomic analyses (**Figure 3**, **Supplemental Tables 6, 7**). The brain, although comprising only 2% of the body mass, has a high metabolic rate and accounts for 20% of the total energy consumption. Consequently, it generates large amounts of ROS (Almeida et al., 2023), which increases under pathological conditions and in aging (Rizor et al., 2019). Astrocytes are the main brain cells responsible for the clearance of ROS: astrocytes express high levels of both non-enzymatic (e.g. glutathione: GSH) and enzymatic (e.g. superoxide dismutase) antioxidant systems (Chen et al., 2020), and astrocytes contain much higher levels of GSH than neurons (Dringen and Arend, 2025). Nuclear factor erythroid 2-related factor 2 (Nrf2), a transcription factor involved in maintaining redox and metabolic homeostasis by regulating the expression of antioxidant enzymes and the biosynthesis of GSH, is also highly expressed by astrocytes, with little expression in neurons (Cuadrado et al., 2019; Dringen and Arend, 2025). Our finding that several antioxidant pathways are activated in CeA astrocytes of dependent mice indicate an adaptive response of reactive astrocytes that compensate for the increased oxidative stress induced by ethanol, although prolonged elevated ROS production may lead to the depletion of astrocyte antioxidant capacity. Under pathological conditions, reactive astrocytes start downregulating their antioxidant functions and instead actively produce ROS (Chen et al., 2020). While most of the changes we identified appear to be increasing the antioxidant capacity of astrocytes, we also observed the activation of pathways (i.e. “production of nitric oxide and reactive oxygen species in macrophages” and “glycation signaling pathway”), that are involved in oxidative stress, suggesting the beginning of a transition toward a maladaptive, pro-oxidant astrocyte phenotype (**Figures 2E**; **3A, B**, **Supplemental Tables 4,6,7**). Future studies testing the oxidative status of the CeA and the inclusion of an alcohol withdrawal time-point is needed for a better understanding of the evolving of pro- and/or antioxidant astrocyte responses in the processes leading to alcohol dependence.

Reactive astrocytes have been shown to have increased branching and surface area, which may affect their function and neuronal support (Baldwin et al., 2023). Using females from the same astro-TRAP transgenic model used in translatome experiments we reconstructed astrocytes and found that major changes in astrocyte morphology occurred in non-dependent mice when compared to naïve. Astrocyte surface area and volume (**Figure 7F,G**) and astrocyte process length and branching, and branch orders (**Figure 8B-G**) were further increased in dependent vs non-dependent animals. Together these findings indicate that changes in astrocyte morphology precede neuroimmune molecular changes associated with astrocyte reactivity and were further enhanced in reactive astrocytes from dependent mice.

Analyses of astro-TRAP-Seq and proteomic datasets of the non-dependent vs naïve comparisons showing overrepresentation of cytoskeleton remodeling pathways support the observation that astrocyte morphological changes occur in the absence of neuroimmune activation (**Figures 2**, **3**; **Supplemental Figures 4, 6, 7**). In addition, major alterations of cytoskeleton remodeling/organization pathways were observed in the phosphoproteomics data sets where we identified “organization of the cytoskeleton” as a significantly regulated function in all comparisons (**Figure 6**). Protein phosphorylation and dephosphorylation are dynamic posttranslational modifications important for the modulation of cytoskeletal plasticity and remodeling (Nixon and Sihag, 1991; Amin et al., 2013; Zhang et al., 2026). Our results indicate that both alcohol drinking and alcohol dependence induce changes in astrocyte morphology via cytoskeleton remodeling.

Together, our findings reveal that alcohol dependence induces molecular and morphological changes in CeA astrocytes that indicate profound astrocyte functional changes consistent with the acquisition of a reactive phenotype. The upregulation of antioxidant and neuroimmune pathways and the downregulation of astrocyte-specific proteins involved in modulating synaptic and blood brain barrier functions suggest that both, adaptive and maladaptive changes occur. In contrast, voluntary alcohol drinking induces changes in astrocytes consistent with maintaining brain homeostasis and with the remodeling of the astrocyte cytoskeleton, that lead to changes in astrocyte morphology. These results are consistent with our recent proteomic studies of cerebrospinal fluid (CSF) of non-dependent and alcohol-dependent mice reporting dependent-specific proteomic signatures of blood-brain barrier disruption, astrocyte-related neuroinflammation, cellular stress responses, and complement system activation (Turner et al., 2026). In contrast, non-dependent specific proteins indicated preserved protective mechanisms including complement regulation, anti-inflammatory signaling, and neuronal calcium homeostasis. This study is a necessary first step toward the identification of astrocyte-specific targets for AUD treatments; the potentiation of reactive astrocyte adaptive changes and the inhibition of their maladaptive alterations represent novel therapeutic approaches to AUD.

## Ethics approval and consent to participate

All procedures were approved by the Scripps Research Institutional Animal Care and Use Committee (IACUC) and are in line with the ARRIVE guidelines and the National Institutes of Health Guide for the Care and Use of Laboratory Animals.

## Acknowledgements

Support for this study was provided by NIH grants: U01AA029965 (MG), P60AA010760 (MG), R01AA029486 (MG), R21DA060442 (MG), U01AA013498 (MR), AA007456 (AG), P60 AA006420 (MR), R37 AA017447 (MR), R01AA021491 (MR), R01AA029841 (MR), K01DA054449 (HN); VA Merit Review Award I01BX001819 (MG); facilities and resources at the Portland VA Health Care System (MG); and the Schimmel Family Chair (MR). RNA sequencing was conducted by the Integrated Genomics Laboratory (RRID: SCR_022651) at OHSU. The contents of this article do not represent the views of the United States Department of Veterans Affairs or the United States government.

---

**Supplemental Figure 1. A.** Confirmation of astrocyte RNA enrichment after TRAP was conducted by qRT-PCR on a subset of IP and input samples using primers for astrocytic (*Aldh1l1*), neuronal (*Tubb3*), microglial (*Itgam*), and oligodendrocyte (*Mbp*) markers. *Aldh1l1* shows enrichment in IP compared to input while the other cell-type markers show de-enrichment in IP samples compared to inputs. **B.** Hierarchical clustering of astro-TRAP-Seq and CeA RNA-Seq samples from Dep, Non-Dep, and Naïve animals from both input (bulk-tissue) and immunoprecipitated (IP) astrocyte translating RNA. The main separation is between IP and input samples with the darker blues indicating IP samples clustering together in the bottom right quadrant of the plot and the input samples clustering together in the upper left quadrant. **C.** Principal component analysis (PCA) of all sequenced samples shows distinct separation of IP and input samples. Samples are colored based on treatment group and fraction. The main separation of samples is along the x-axis (PC1), separating input and IP samples. **D.** PCA of only the IP samples only shows little separation of Dep (D), Non-Dep (ND), and Naïve (N) samples. **E.** PCA plot showing the sex of each sample with females in red and males in blue. **F.** PCA plot showing individual TRAP passes with group E showing some separation from the other TRAP passes.

**Supplemental Figure 2.** Summary of astro-TRAP-Seq and input RNA-Seq from three comparisons of CIE-2BC treatment groups. **A.** Venn diagram showing overlap of Dep (D) vs. Naïve (N), Dep (D) vs Non-Dep (ND), and Non-Dep (ND) vs Naïve (N) treatment group comparisons of astro-TRAP-Seq DT genes. **B.** Venn diagram showing overlap of Dep (D) vs. Naïve (N), Dep (D) vs Non-Dep (ND), and Non-Dep (ND) vs Naïve (N) treatment group comparisons of input DE genes.

**Supplemental Figure 3**. Pathway analysis of CeA proteomics. **A.** Top 30 IPA canonical pathways from the Dep vs. Naïve comparison. **B.** Top 30 IPA canonical pathways from the Dep vs. Naïve comparison with regulated proteins minus regulated proteins whose genes are low expressed in astrocytes compared to the bulk tissue. **C.** List of overlapping changes in expression of genes (by astro-TRAP-Seq) and proteins (by proteomic analysis) from the Non-Dep vs. Naïve comparison. 15 genes were regulated in the astrocyte TRAP dataset were also regulated in the proteomics analysis, including 10 proteins enriched in astrocytes; 13 upregulated, 8 involved in immune functions and 1 protein regulated in opposite direction than its RNA in TRAP-Seq. In panels **A-B**, the x-axis shows the activation z-score which is a prediction of overall pathway activation or inhibition based on known pathway relationships and the regulated proteins in the pathway, the size of each circle represents the proportion (ratio) of significant proteins to the total pathway size, and the color of each circle corresponds to the enrichment p-value, with brighter colors corresponding to lower p-values. Pathway names related to immune responses are highlighted in yellow, oxidative stress/antioxidant pathways are highlighted in blue, and cholesterol-related pathways are highlighted in orange.

**Supplemental Figure 4.** IPA canonical pathway analysis of differentially phosphorylated proteins in CeA after CIE-2BC. Proteins with significantly altered phosphorylation were used to determine the top 30 canonical pathways in the Dep vs Naïve (**A**), Dep vs. Non-Dep (**B**), and Non-Dep vs. Naïve (**C**) comparisons. Proteins whose expression was regulated in the same direction as the phosphorylation changes were removed from these analyses. In all panels, the x-axis shows the activation z-score which is a prediction of overall pathway activation or inhibition based on known pathway relationship and the regulated proteins in the pathways, the size of each circle represents the proportion (ratio) of significant phosphorylated proteins to the total pathway size, and the color of each circle corresponds to the enrichment p-value, with brighter colors corresponding to lower p-values. Pathways highlighted in green are related to RHO/cytoskeleton remodeling, yellow are related to neuroinflammation, and blue are related to oxidative stress.

