## Supplemental Methods for "Astrocyte Reactivity by Alcohol Dependence in the Central Amygdala"

### Supplementary Methods

#### CIE-2BC Detailed Methods

CIE-2BC detailed methods were conducted as described (Hashimoto et al., 2025a). Following acclimation, mice were singly housed and given two hours of access to two drinking tubes, one containing 15% ethanol and the other containing water (i.e. two bottle choice or 2BC) for 20 days (5 days per week for 4 weeks). Consumption of ethanol and water was recorded during these 2-hour periods. After the baseline period, animals were split into two balanced groups based on equal ethanol consumptions. Mice were exposed in their home cages, with Dependent animals receiving intermittent ethanol vapor and Non-Dependent mice receiving control air. The Dependent cohort was injected with ethanol + pyrazole (alcohol dehydrogenase inhibitor, see below for details) and cages were moved to the chambers to receive intermittent vapor exposure for 4 days (16hr vapor on, 8hr off). Each Thursday, directly following 16hrs of exposure to vapor, cages were removed, and tail blood was collected for blood ethanol concentration (BEC) determination. Target BEC range was175-200 mg/dl. Following the fourth day of exposure, mice were left undisturbed for 72 hours and were then once again given the two-bottle choice (5 days of 2-hour access) to measure ethanol preference and consumption following vapor chamber exposure. At the same time as the vapor groups, the Non-Dependent mice were injected with pyrazole in saline and received 2BC testing. The vapor/air exposure and 5 days of 2 bottle choice testing were repeated 3 more times for a total of 4 rounds of exposure and 2BC testing. A final (5^th^) round of CIE exposure was completed, and mice were euthanized for whole-brain collection within an hour of removal from the chambers. Brains were flash frozen in isopentane chilled in a dry ice isopropanol slurry and stored at -80°C until processing. Additional whole-brains were collected and processed from an age-matched ethanol-naïve astrocyte-TRAP cohort which was left undisturbed in their home cages (i.e. treatment naïve).

#### Ethanol vapor exposure

Ethanol (95%) was pumped into a 2000 ml Erlenmeyer vacuum on a warming tray (50°C) to create ethanol vapor. Ethanol vapor was introduced independently into each sealed chamber (Quad Passive System, La Jolla Alcohol Research, Inc., La Jolla, CA) via a stainless-steel manifold (11 L/min). Throughout exposures, mice were kept in their home cages. Ethanol vapor concentrations were manipulated by varying the ethanol flow rate into the flask and typically ranged from 22 to 27 mg/liter. To more finely control blood alcohol levels, the pyrazole and ethanol loading dose were varied between 0.875-1.75 g/kg (ethanol) and 34-68.1 g/kg (pyrazole). Both the Dependent and Non-Dependent mice cohorts were given the same pyrazole dosage. Monitoring occurred at least once daily while mice were in the vapor chambers and bloods were taken from dependent animals weekly to allow for optimal vapor chamber regulation as well as to assess each mouse’s health.

#### Blood alcohol concentration for dependent mice

Blood samples were obtained by cutting 0.5 mm from the tip of each mouse's tail with a clean surgical blade. Approximately 40 µl blood was collected using capillary tubes and then emptied into Eppendorf tubes containing evaporated heparin and kept on ice. Samples were centrifuged, and plasma collected into fresh Eppendorf tubes. 5 µl plasma was injected into an Agilent 7820A GC coupled to a 7697A (headspace-flame-ionization). Results were calibrated and compared with a 6-point serial diluted calibration curve of 300 mg/dl ethanol (Cerilliant E-033).

#### Translating Ribosome Affinity Purification

Frozen brain punches were homogenized in homogenization buffer using a disposable pellet pestle (Kontes) and astrocyte translating RNA was isolated using the TRAP procedure as described previously (Hashimoto et al., 2025b), using anti-GFP antibodies from the Memorial-Sloan Kettering Antibody and Bioresource Core Facility (HtzGFP-19C8 & HtzGFP-19F7) and protein A/G magnetic beads (Thermo Fisher, 88803). RNA was isolated from the input and IP samples using Trizol (Thermo Fisher, Carlsbad, CA) and the Direct-Zol RNA MicroPrep Kit (Zymo Research, Orange, CA).

#### RNA-Seq data processing and analysis

Sequencing reads were quality assessed with FastQC v0.11.9 (Andrews, 2010), followed by trimming with Trimmomatic v0.39 (Bolger et al., 2014) and alignment to the mouse reference genome from Ensembl (GRCm38) with the STAR aligner v2.7.10b (Dobin et al., 2013). The Ensembl annotation gtf file GRCm38.99 was used during alignment to define gene regions and to obtain gene counts from the ReadsPerGene output files from STAR. To determine potential bias in EGFP expression, reads were aligned to the EGFP gene using bwa-mem v0.7.17 (Li, 2013) and EGFP expression was determined by taking the number of read pairs mapped to EGFP divided by the input read pairs aligned to EGFP, multiplied by one million. Genes with low counts were filtered out of each dataset (RNAseq, TRAPseq, combined), using the filterByExpr function from edgeR v3.28.0 (Robinson and Oshlack, 2010).

#### Proteomics & Phosphoproteomics Detailed Methods

*Sample preparation*

A solution of 8M urea (MilliporeSigma, St. Louis, MO) prepared in 50 mM NH_4_HCO_3_ (Sigma-Aldrich) was added to the mouse brain punch samples. Next, the samples were homogenized with a pellet pestle (VWR, Radnor, PA) and protein amount was determined by Pierce BCA Protein Assay reagents (ThermoFisher Scientific, Waltham, MA). A 100 mM Dithiothreitol (DTT) (MilliporeSigma) solution was added to reach a final concentration of 10 mM, followed by incubation at 37 °C for 60 min with 850 rpm shaker speed. A 400 mM iodoacetamide (MilliporeSigma) solution was subsequently added to each sample with a final concentration of 40 mM, followed by incubation at 37 °C for 60 min with 850 rpm shaker speed in dark. Next, samples were diluted by 8-fold using 50 mM NH_4_HCO_3_ and 1 M CaCl_2_ (MilliporeSigma) was added for a final concentration of 1 mM CaCl2. Trypsin (Promega Corporation, Madison, WI) was warmed to 37 °C for 5-10 minutes prior to being added to samples at 1:50 enzyme:protein ratio. The samples were incubated at 37 °C for 6 hours with shaker speed at 850 rpm. To perform SPE peptide cleanup, C-18 cartridges (Phenomenex, Torrance, CA) were pre-conditioned with 3 mL of 100% MeOH (Fisher Scientific, Waltham, MA) using a vacuum manifold (Supelco, Bellefonte, PA), followed by 1 mL of 0.1% TFA (MilliporeSigma). The digested samples were then loaded onto the pre-conditioned SPE cartridges. The cartridges were washed using 4 mL of 95:5 H2O:ACN (Fisher Scientific) and peptides were eluted from the SPE cartridges using 1 mL of 80:20 ACN:H2O. The peptides were concentrated in a centrifugal evaporator (Thermo Savant, Waltham, MA) before being quantified using Pierce BCA Protin Assay reagents. Peptides were dried down in a speed-vac and resuspended in 3% ACN, 0.1% formic acid (MilliporeSigma) with a final concentration of 0.1 µg/µl.

For phoshoproteomics analysis, Fe^3+^-NTA agarose beads were freshly prepared using the Ni-NTA Superflow agarose beads (cat. no. 30410; QIAGEN) to enrich phosphopeptides as previously described (Hardesty et al., 2022). For each AMY sample, the remaining peptides from global proteomics analysis were reconstituted to 0.5 µg/µL in IMAC binding/wash buffer (80% acetonitrile, 0.1% trifluoroacetic acid) and incubated with 10 μL of the Fe^3+^-NTA agarose beads for 30 min at room temperature. After incubation, the beads were washed 2 times each with 50 μL of wash buffer and once with 50 μL of 1% formic acid on the stage tip packed with 2 discs of Empore C18 material (Empore Octadecyl C18, 47 mm; cat. no. 98-0604-0217-3; CDS Analytical). Phosphopeptides were eluted from the beads on C18 using 70 μl of elution buffer (500 mM potassium phosphate buffer); 50% acetonitrile and 0.1% formic acid were used for the elution of phosphopeptides from the C18 stage tips. Samples were dried using Speed-Vac and later reconstituted with 12μL of 3% acetonitrile and 0.1% formic acid containing 0.01% n-dodecyl-beta-maltoside for LC–MS/MS analysis.

#### *Mass spectrometry data acquisition*

Peptides and phosphopeptides were analyzed on Orbitrap Astral mass spectrometer (Thermo Scientific), acquiring full scan spectra by Orbitrap analyzer with a scan range of 380 to 980 m/z at the resolution of 240,000. The normalized AGC target was set at 500% with maximum injection time at 5 ms and the RF lens was set at 45%. DIA window type was set to “Auto”, window placement optimization at “On”, window overlap at 0, and isolation window at 2 m/z. Higher-energy collisional dissociation was performed at a normalized collision energy of 25% and loop control was set at 0.6 seconds.

Liquid chromatography separation was performed using a Vanquish Neo LC (Thermo Scientific) with a 70 SPD (Samples Per Day) separation method. Each sample had a 14-minute active gradient along with 6-minute sample loading and column equilibration. The Vanquish Neo was configured to the trap-and-elute mode utilizing the PepMap Neo Trap Cartridge (Thermo Scientific). A PepMap ES906 analytical column was used for reverse phase peptide elution. The analytical column was interfaced to the mass spectrometer using an EASY-Spray source. The ion source conditions were set to 2.2 kV for the spray voltage and 300 °C for ion transfer tube temperature. Mobile phases consisted of (A) 0.1% formic acid in water and (B) 0.1% formic acid in acetonitrile with the following gradient profile (min, Flow Rate, %B): 0.0, 3.50,1.0; 0 .001, 2.80, 1.0; 0 .501, 1.30, 4.0; 0 .901, 1.10, 8.0; 12.301, 1.10, 22.5; 15.301, 1.10, 35.0; 15.501, 1.10, 55.0; Column Wash; 16.001, 2.80, 99.0; 17.001, 2.80, 99.0; 18.000, 2.80, 1.0.

#### *Proteomics data processing and statistical analysis*

The mass spectrometry raw files were processed in a batch mode and searched together using DIA-NN (Demichev et al., 2020) (Version 1.8.1) against UniProt fasta of mouse proteome (2023-03-01-reviewed with contaminants, 21,949 entries) in default parameters with several modifications. In silico digest was activated with cleavage at K, R residues to keep a maximum number of missed cleavages set to 2. Peptide length was set 7–30 with a precursor m/z range of 380–980 and charge state +2 to +4. Variable modifications in global proteomics search included mass delta 15.9949 for oxidation at Met and mass delta 42.0106 for protein N-terminal acetylation. For phosphoproteomic search, serine/threonine/tyrosine phosphorylation (mass delta 79.9663) was added to the variable modification list applied in global proteomics search. Cys alkylation with mass delta 57.02146 was set as static modification. Precursor ion generation was activated for both FASTA digest and deep learning-based spectra. Protein inference was set to gene level with single-pass mode neural network classifier, and RT-dependent cross-run normalization, MBR activation, and 1%FDR filtering.

#### Astrocyte Morphometric Analysis

The Aldh1l1-EGFP ribosomal fluorescent tag was amplified in the CeA using a free-floating protocol. Sections were incubated for 1hr in 5% normal goat serum (NGS, Sigma-Aldrich) and 0.25% Triton-X (Sigma-Aldrich) in 1-X PBS. Tissue was then incubated for 72hr at room temperature in 5% NGS 0.25% Triton-X and the primary antibody;1:800 rat anti-EGFP (Nacalai Cat#: 04404-84, RRID: AB_10013361). Sections were rinsed in 1-X PBS and incubated in 5% NGS, 0.25 % Triton-X and the secondary antibody; 1:500 Alexa Fluor donkey anti-Rat 488 (Thermo Fisher Scientific, Cat#: A-21208, RRID: AB_2535794) for 4hr at room temperature. Tissue was then rinsed in 1-X PBS, incubated for 10 minutes in 1:5,000 DAPI (Anaspec Cat#: AS-83210) in 1-X PBS, and rinsed again in 1-X PBS. Sections were mounted into FrostPlus slides and cover slipped for imaging.

*Microscopy and astrocyte 3D reconstruction*

The CeA was imaged using a Zeiss LSM 780 Confocal microscope (Zeiss, Oberkochen, Germany). Standardization of laser intensity, gain, offset, and pinhole settings were determined using a tissue section containing the CeA from a naïve control animal. The same settings were maintained for all subsequent images. Four bilateral images were captured per animal with a 63X (1.40 numerical aperture) oil immersion lens. Z-stacks ranged between 49-53μm with a z-step 0.38μm. Images (0.085 pixels/μm) were then exported as .tif files.

Images were converted from .tif files to .jpx files using MicroFile+ (V2023.1.3; MBF Bioscience, Williston, VT). Images were imported into the Neurolucida 360 environment (V2024.2.2; MBF Bioscience, Williston, VT) and the image background intensity was corrected using a rolling ball radius of 15. Three-dimensional (3D) reconstructions were drawn first by identifying the cell body of the astrocyte, which was traced manually throughout the volume of the image stack, setting the user-guided mode to trace “trees” and “bifurcating trees” of the astrocytes. Manual tracing was used at points where semi-automated software detection was not successful in identifying branches in user-guided mode. Approximately 6-9 reconstructions were completed per animal (1-4 reconstructions per image). Once reconstructions were completed, the data were exported to Neurolucida Explorer (V2022.2.1; MBF Bioscience, Williston, VT) to generate morphometric analyses and quantitative measurements of the astrocytic reconstructions. These analyses included process branch length, the number of processes, the number of nodes, complexity index ([Sum of the terminal orders + Number of terminals]* [Total dendritic length / Number of primary dendrites]), Convex hull 2D and 3D (dendritic field size), intersections, endings, branch complexity analyses, and Sholl analyses. The sample size for astrocyte morphological assessments was a total of 19-20 cells from 3 animals per treatment group.
