## Supplemental Figures for "Astrocyte Reactivity by Alcohol Dependence in the Central Amygdala"

### Supplemental Figure 1

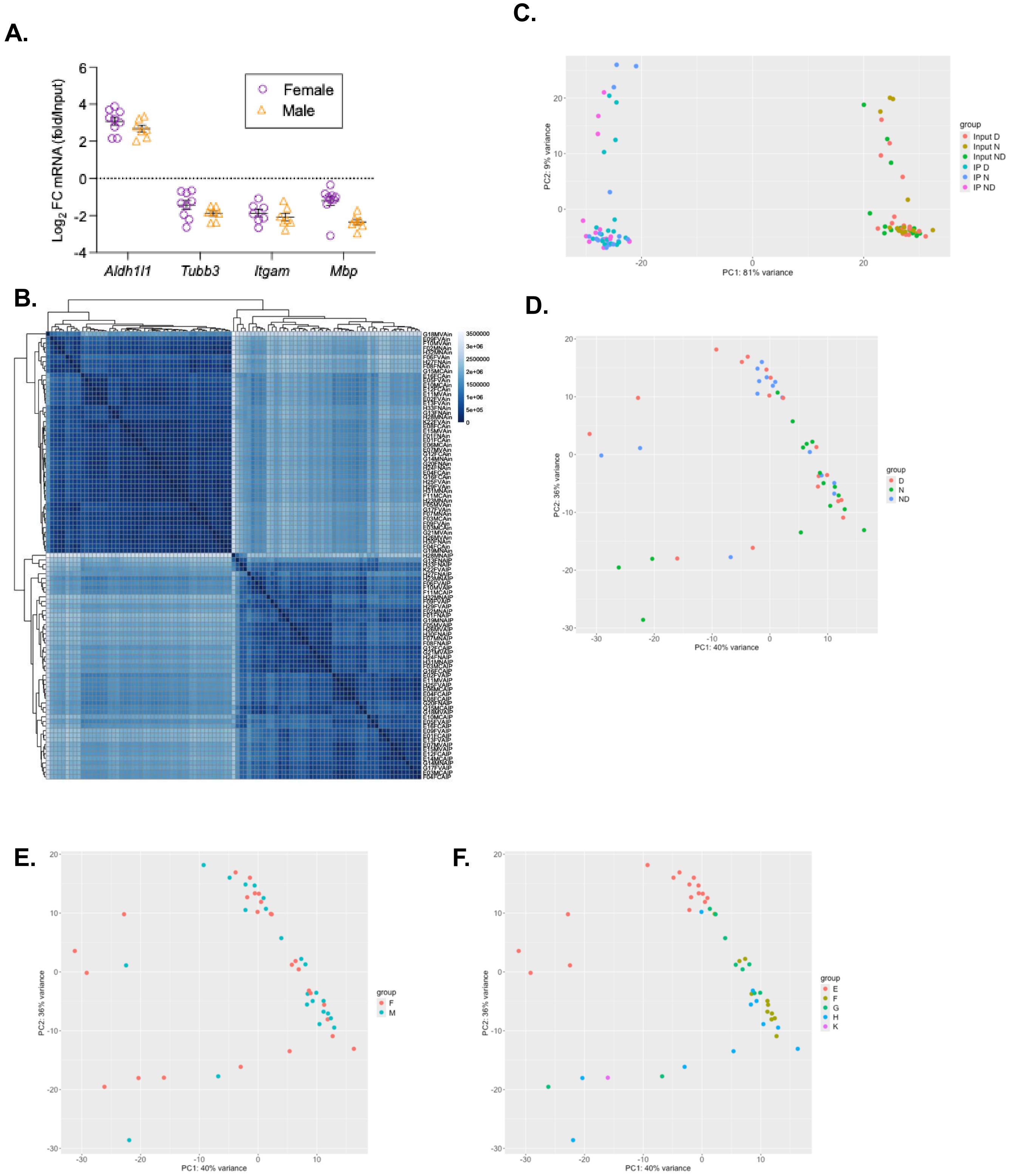

### Supplemental Figure 2

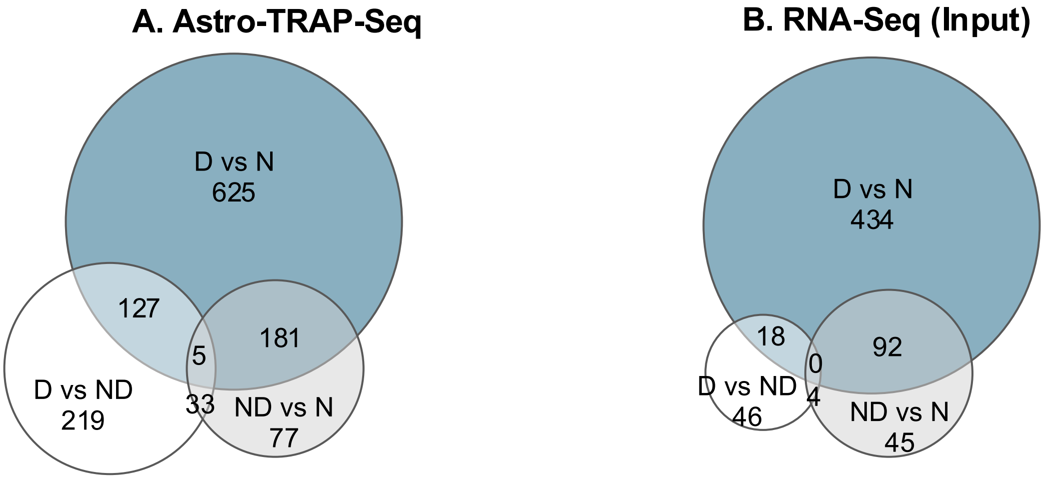

### Supplemental Figure 3

**A.** Dep vs. Naive

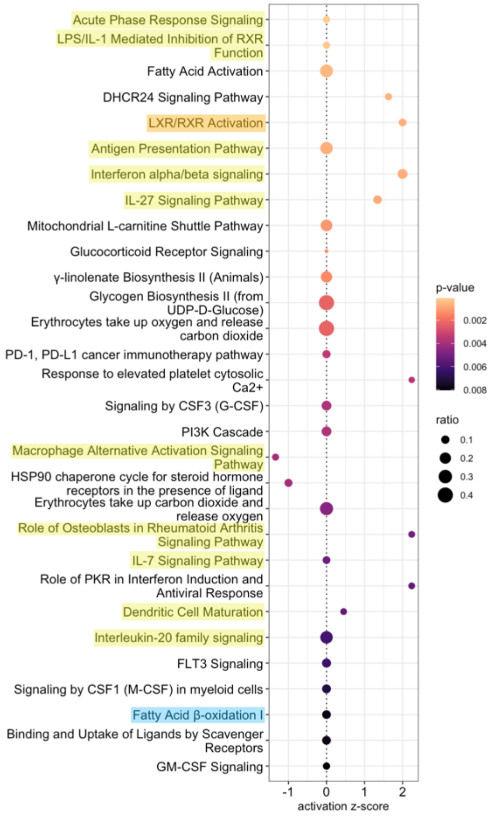


**B.** Dep vs. Naive (Minus proteins less expressed in astrocytes)

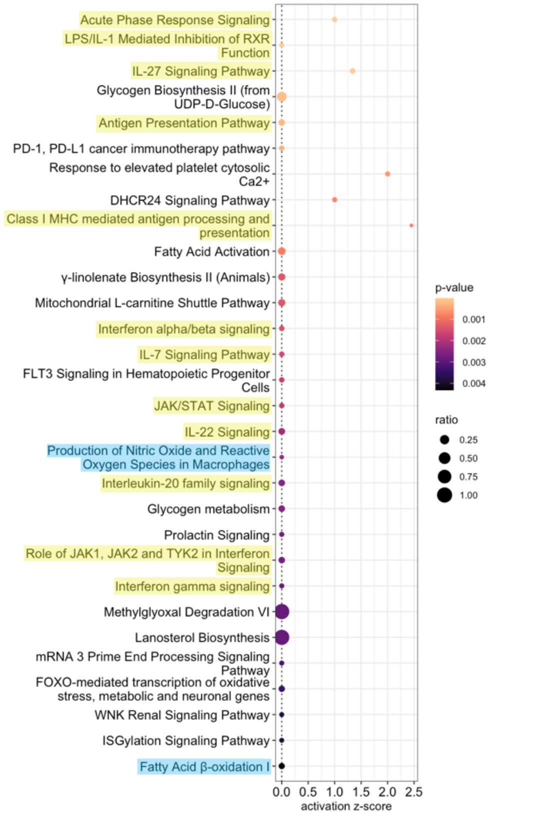


**C.**

| Gene Symbol | Name | Proteomics | | | TRAP RNA-Seq | | |
|---|---|---|---|---|---|---|---|
| | | logFC | P.Value | adj.P.Val | log2FC | pvalue | padj |
| **Aass** | **aminoadipate-semialdehyde synthase** | **0.4170** | **6.39E-07** | **2.26E-03** | **0.4668** | **1.48E-05** | **1.11E-03** |
| Cspg4 | chondroitin sulfate proteoglycan 4 | 0.2436 | 2.09E-05 | 1.39E-02 | 0.4600 | 2.26E-03 | 4.35E-02 |
| **Lgals3bp** | **lectin, galactoside-binding, soluble, 3 binding protein** | **0.9971** | **3.51E-05** | **1.85E-02** | **0.7983** | **1.22E-03** | **2.90E-02** |
| **Plin4** | **perilipin 4** | **0.7409** | **6.78E-05** | **2.37E-02** | **1.6750** | **1.06E-05** | **8.45E-04** |
| ***Pik3ip1*** | ***phosphoinositide-3-kinase interacting protein 1*** | ***0.3719*** | ***9.38E-05*** | ***2.79E-02*** | ***-0.4704*** | ***3.53E-08*** | ***7.64E-06*** |
| Hspa8 | heat shock protein 8 | -0.1237 | 1.64E-04 | 3.18E-02 | -0.4717 | 3.35E-09 | 1.08E-06 |
| Ctss | cathepsin S | 0.5382 | 2.71E-04 | 3.99E-02 | 0.9326 | 7.43E-04 | 2.08E-02 |
| Fkbp5 | FK506 binding protein 5 | 0.2256 | 3.91E-04 | 4.56E-02 | 0.7401 | 1.12E-05 | 8.94E-04 |
| **Irgm1** | **immunity-related GTPase family M member 1** | **1.0472** | **6.05E-04** | **5.53E-02** | **1.3691** | **6.07E-04** | **1.80E-02** |
| **Igtp** | **interferon gamma induced GTPase** | **1.9465** | **7.17E-04** | **6.03E-02** | **2.3298** | **7.08E-06** | **6.26E-04** |
| **Acadsb** | **acyl-Coenzyme A dehydrogenase, short/branched chain** | **0.1152** | **8.74E-04** | **6.54E-02** | **0.2400** | **2.64E-03** | **4.79E-02** |
| **Stat1** | **signal transducer and activator of transcription 1** | **1.0961** | **1.22E-03** | **7.39E-02** | **0.9624** | **1.88E-03** | **3.83E-02** |
| **Gbp2** | **guanylate binding protein 2** | **1.3653** | **1.58E-03** | **8.37E-02** | **2.1465** | **3.86E-05** | **2.34E-03** |
| Hnrnpc | heterogeneous nuclear ribonucleoprotein C | -0.1284 | 1.68E-03 | 8.42E-02 | -0.4282 | 1.08E-07 | 2.08E-05 |
| **B2m** | **beta-2 microglobulin** | **0.8263** | **1.92E-03** | **8.99E-02** | **0.8950** | **2.85E-03** | **5.00E-02** |

**Bold text indicates gene is enriched in astrocytes in the CeA (TRAP IP vs Input padj < 0.05 & log2FC > 1)**

*Italics* text indicates opposite direction of regulation between TRAP-Seq and proteomics

Underline indicates gene involved in immune pathwyas

### Supplemental Figure 4

**A.** Phosphoproteomics: Dep vs. Naive

**B.** Phosphoproteomics: Dep vs. Non-Dep

**C.** Phosphoproteomics: Non-Dep vs Naive

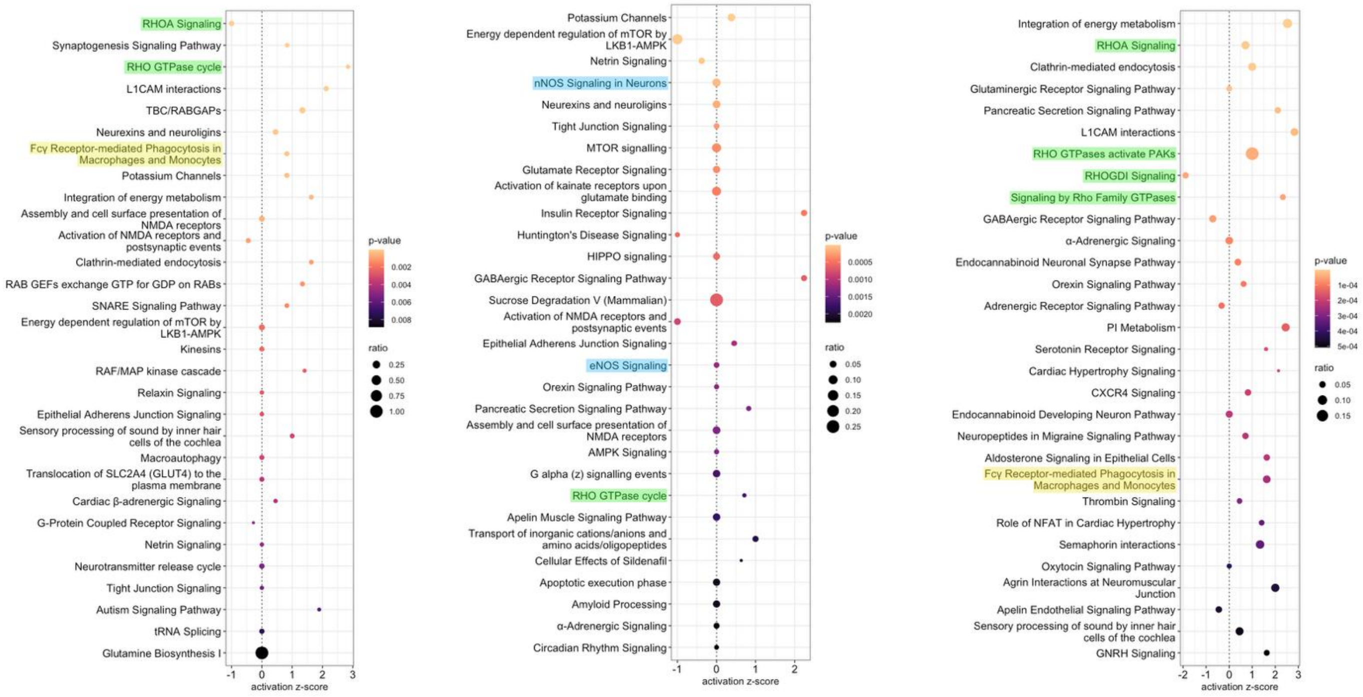
